# Temporal progression along discrete coding states during decision-making in the mouse gustatory cortex

**DOI:** 10.1101/2022.07.20.500889

**Authors:** Liam Lang, Giancarlo La Camera, Alfredo Fontanini

## Abstract

The mouse gustatory cortex (GC) is involved in taste-guided decision-making in addition to sensory processing. Rodent GC exhibits metastable neural dynamics during ongoing and stimulus-evoked activity, but how these dynamics evolve in the context of a taste-based decision-making task remains unclear. Here we employ analytical and modeling approaches to i) extract metastable dynamics in ensemble spiking activity recorded from the GC of mice performing a perceptual decision-making task; ii) investigate the computational mechanisms underlying GC metastability in this task; and iii) establish a relationship between GC dynamics and behavioral performance. Our results show that activity in GC during perceptual decision-making is metastable and that this metastability may serve as a substrate for sequentially encoding sensory, abstract cue, and decision information over time. Perturbations of the model’s metastable dynamics indicate that boosting inhibition in different coding epochs differentially impacts network performance, explaining a counterintuitive effect of GC optogenetic silencing on mouse behavior.

## Introduction

Over the past decade, the gustatory cortex (GC) has emerged as a model for studying cortical dynamics. Analysis of neural activity in alert rats has revealed that ensembles of GC neurons consistently hop between semi-stable patterns of activity (Jones et al., 2007; Moran and Katz, 2014; Mazzucato et al., 2016; Mazzucato et al., 2015; Sadacca et al., 2016) termed “metastable dynamics” (for reviews, see La Camera et al., 2019; Brinkman et al., 2022). While these metastable dynamics were originally found in the context of taste processing, recent work has reported that metastability can also be associated with cognitive processes such as taste expectation (Samuelsen et al., 2012; Mazzucato et al., 2019) and sensorimotor transformations (Sadacca et al., 2016; Mukherjee et al., 2019). The latter was shown in the case of rats producing gapes—orofacial reactions signaling aversion and rejection (Grill and Norgren, 1978)—in response to bitter stimulation. During the course of such a consummatory experience, GC neural ensembles switch from taste-encoding to gape-anticipating metastable states (Sadacca et al., 2016; Mukherjee et al., 2019). Optogenetic manipulations of activity preceding this switch lead to a delayed onset of gapes (Mukherjee et al., 2019), demonstrating a causal role of GC’s metastable dynamics in naturalistic consummatory behavior.

Recent studies indicated that, in addition to representing taste and naturalistic sensorimotor transformations, GC can also contribute to taste-based decisions in mice engaged in a decision-making paradigm (Vincis et al., 2020). Mice were given one of four tastants—two sweets and two bitters—and, after a brief delay period, had to register a decision by licking one of two lateral spouts. Electrophysiological recordings in mice performing this task uncovered that GC neurons encode sensory and decision-making information during the sampling and delay periods. Interestingly, and counterintuitively for a sensory region, optogenetic perturbation of GC activity during stimulus sampling had no impact on behavioral performance, whereas silencing during the delay period significantly impaired task performance (Vincis et al., 2020). While important in highlighting the role of GC in taste-based decision-making, this research raised two fundamental questions: 1) Is GC’s activity metastable during decision-making and, if so, what role does GC metastability play in perceptual decisions? 2) Why is optogenetic perturbation of GC neural activity during stimulus sampling less disruptive than during the delay period?

Here, we investigate these questions using a Hidden Markov Model (HMM) analysis of the data (Radons et al., 1994; Abeles et al., 1995; Jones et al., 2007; Ponce-Alvarez et al., 2012; Mazzucato et al., 2015; Sadacca et al., 2016) and a biophysically grounded model of GC neural activity. Using HMM, we demonstrate the existence of a temporal progression of metastable states emerging, sequentially, for coding sensory information (taste quality), internal deliberations, and finally forthcoming action. To understand the computational mechanisms underlying the genesis of metastable activity and the emergence of coding states in the context of decision-making, we developed a metastable spiking network model of GC. The model provides a mechanistic understanding of the results of our HMM analysis—in particular, the temporal progression of hidden coding states for different sensory and cognitive variables. The model is then used to explain the experimental effects of the optogenetic perturbation—in particular, why performance is more sensitive to perturbations during the delay period than during the stimulus-evoked period (Vincis et al., 2020)—and investigate the effects of perturbations on metastable dynamics.

Altogether, our results show that GC supports complex taste-related decisions via sequences of metastable states that follow a precise temporal progression. The mechanism for this progression is explained by a biologically plausible model of GC, which also clarifies the counterintuitive effects of optogenetic silencing on performance, and further provides testable predictions on the effects of intensity and timing of optogenetic perturbations on metastability and behavioral performance.

## Results

### Decision-making neural activity in mouse GC is metastable

Mice implanted with drivable multi-tetrode arrays in the gustatory cortex were trained on a taste-based perceptual decision-making task (Vincis et al., 2020). In this task, head-fixed, water-restricted animals sampled a tastant—sucrose, quinine, maltose, or sucrose octaacetate—from a central spout and learned to lick one of two lateral spouts based on the following rule: sucrose or quinine means lick left, and maltose or octaacetate means lick right. A correct directional lick resulted in a water reward from the lateral spout. The intentional counterbalancing of taste quality between cued directions—sucrose and maltose are sweet, while quinine and octaacetate are bitter—ensured that the animal could not rely simply on the generalization of sweet or bitter to perform well on the task, but rather had to recategorize the four stimuli according to the rule. A variable delay between sampling and availability of the lateral spouts was implemented such that the average time between the sampling/taste event (first lick to the central spout) and the decision event (first lick to the lateral spout) was 2.53 s (range: 1.40 − 6.66 s) (**Figure 1a**). Once mice learned the task (i.e., they reached criterion performance of at least 75% for 3 consecutive days), recording sessions began. We recorded from a total of 16 sessions in four mice, with an average of 198 trials per session (range: 75 − 292) and 5 simultaneously recorded neurons per session (range: 3 − 9).

**Figure 1.**
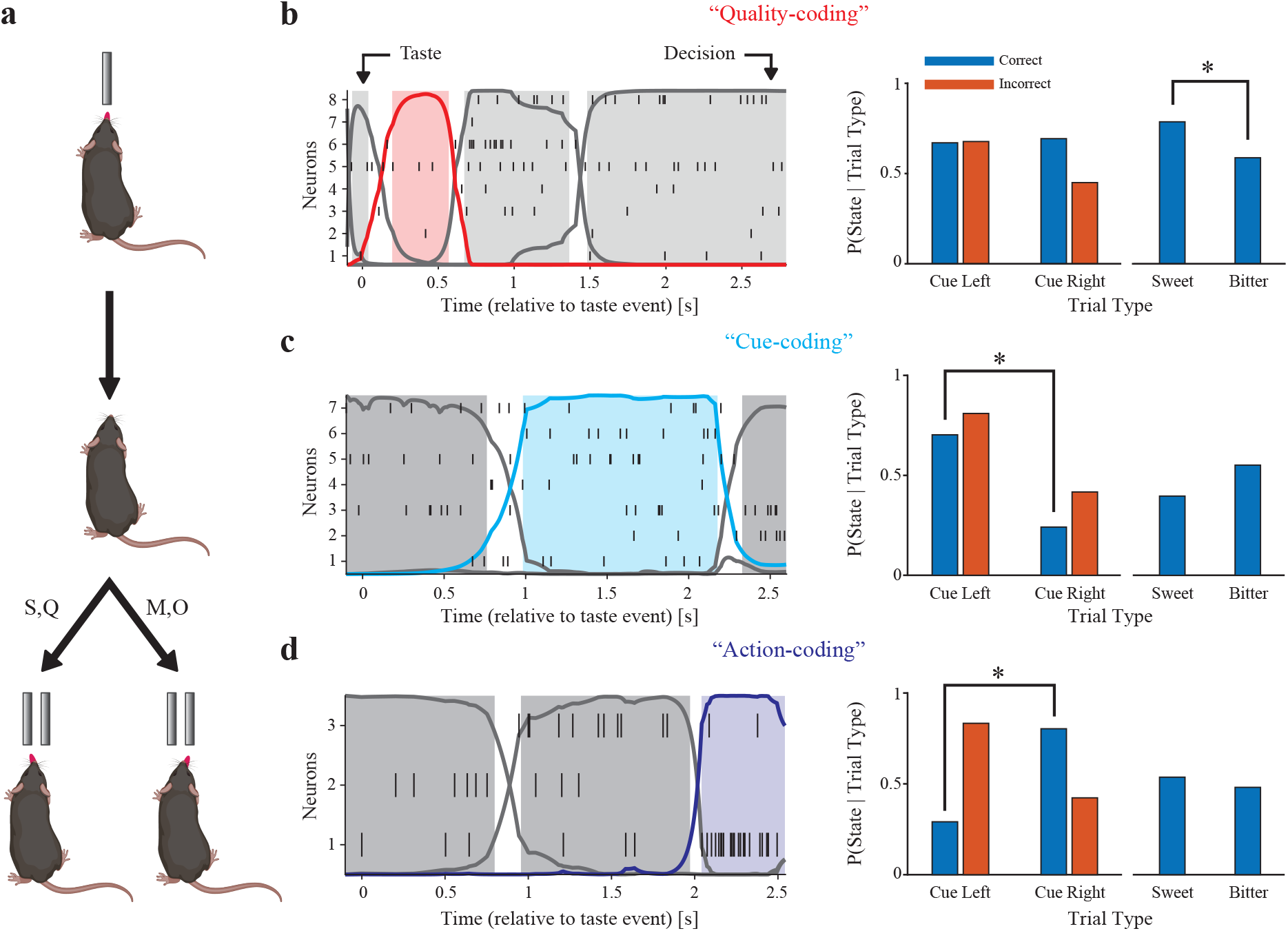
Decision-making activity in GC is metastable and distinct metastable states encode different task-relevant variables. **a**: Perceptual decision-making task schematic (Vincis et al., 2020). S: sucrose, Q: quinine, M: maltose, and O: sucrose octaacetate. **b-d (left)**: Raster plots and HMM-decoded metastable states for in vivo spiking activity during decision-making behavior. “Taste” indicates the animal’s first lick to the central spout; “Decision” indicates the animal’s first lick to a lateral spout. **b-d (right)**: Classification of hidden states. Quality-coding (**b**, red), Cue-coding (**c**, cyan), and Action-coding (**d**, blue) states. * indicates significant difference by Chi-squared test (*p* < 0.05, corrected for the number of decoded states in the session).

We hypothesized that metastable dynamics support the ability of GC to code for different task-related variables, informed by recent findings revealing a crucial role for GC in sensory and decision-making aspects of this task (Vincis et al., 2020), as well as work demonstrating metastability in GC (Jones et al., 2007; Mazzucato et al., 2015). To test this hypothesis, we fitted a Hidden Markov Model (HMM) to our ensembles of simultaneously recorded spike trains. This method allowed us to segment the neural activity into recurrent sequences of discrete states that are not directly observable—a signature of metastable dynamics previously described in this area (Jones et al., 2007; La Camera et al., 2019; Brinkman et al., 2022). The hidden states were defined as vectors of firing rates across the simultaneously recorded neurons (Gat and Tishby, 1993; Seidemann et al., 1996; Jones et al., 2007; Bollimunta et al., 2012; Ponce-Alvarez et al., 2012; Mazzucato et al., 2015).

HMM was fitted to the spike trains of each session separately, and then used to decode, trial-by-trial, the sequence of hidden states (see **Methods** for details). The neural activity in all sessions and trial types could be characterized by sequences of metastable states, with a median of 4.5 hidden states per session (range: 3 − 8). Examples of state sequences are shown in **Figure 1b-d (left panel)**. Note how state transitions occur at random times in different trials. The distribution of state durations resembled an exponential distribution with mean 577 ms (standard deviation: 602 ms; median: 338 ms; range: 50 − 5, 546 ms), in keeping with previous findings in the rat GC (Jones et al., 2007; Mazzucato et al., 2015). These results demonstrate that ensemble activity of mouse GC during decision-making is metastable. Remarkably, some states occurred more often in specific trial types, with their onset times obeying distinct probability distributions, as we show next.

### Distinct discrete states encode task-related variables

To identify whether hidden states encoded task-related variables, state occurrences were compared across different trial types (i.e., characterized by different stimuli or actions; see **Figure 1b-d, right** and **Supplementary Figure 1**). For a state to be considered a coding state, occurrence frequencies had to differ across correct trials of the four different stimuli (Chi-squared test, *p* < 0.05). An additional comparison was performed to determine whether the probability of observing the state was significantly different for one stimulus out of the four (assessed by running pairwise Marascuilo post-hoc tests; see **Methods**). If this was the case, the state was considered “Taste ID-coding.” If not, we grouped correct trials by cued direction (cue left: sucrose or quinine; cue right: maltose or octaacetate) and by taste quality (sweet: sucrose or maltose; bitter: quinine or octaacetate). If the state appeared with significantly different frequency between correct cue left and correct cue right trials, but not between correct sweet and correct bitter trials, it was considered a “Decision-coding” state. Similarly, if a state appeared with significantly different frequency between correct sweet and correct bitter trials, but not between correct cue left and correct cue right trials, it was considered a “Quality-coding” state. In the case that both of these comparisons were significantly different, the state was considered “Dual-coding” (see **Methods** for details). Over all the 16 recording sessions we found 8 Taste ID-coding states, 3 Dual-coding states, 15 Decision-coding states, and 4 Quality-coding states. We chose to focus on Quality-coding and Decision-coding states because they have intuitive interpretations in the context of our behavioral task: Quality-coding states are associated with either sweet or bitter taste qualities, which reflect processing of sensory information. Decision-coding states are associated with either the left or right directions, which reflect decision-making activity. These states were distributed sparsely across sessions: 62.5% of sessions contained a Decision-coding state and 25% of sessions contained a Quality-coding state, with only 18.8% of sessions containing both (**Supplementary Table 1, Column 1**).

Decision-coding states were additionally divided into two categories: “Cue-coding” and “Action-coding” states, the former being associated with the perceptual categorization of the stimulus based on the direction it cues (i.e., sucrose or quinine → left; maltose or octaacetate → right), and the latter being associated with the execution of the action (i.e., go left; go right). This separation was based on the frequency of occurrence of states in correct and incorrect trials. Decision-coding states associated with the same cued direction in correct and incorrect trials were deemed Cue-coding since they were consistently associated with the cued direction, whereas Decision-coding states associated with opposite cued directions in correct and incorrect trials were deemed Action-coding since they were consistently associated with the chosen direction (see **Methods** for details). We found 7 Action-coding states and 8 Cue-coding states (**Supplementary Table 1, Column 1**). Imposing stricter criteria at this step of the classification (i.e., requiring that sessions contain at least 10 incorrectly performed trials of each cued direction) did not qualitatively alter any of the results shown here, but decreased the number of Action- and Cue-coding states to 5 and 6, respectively.

### Temporal sequences of coding states reflect task structure

Previous literature highlights the importance of temporal dynamics in GC (Katz et al., 2001; Fontanini and Katz, 2006; Jones et al., 2007; Sadacca et al., 2012; Sadacca et al., 2016; Mazzucato et al., 2019), with different variables being encoded at different times following stimulus delivery. To determine whether the variables associated with a taste-based, 2-alternative choice (2-AC) task are encoded sequentially, we analyzed the within-trial onset times of coding states. Consistent with the literature, we hypothesized that Quality-coding states should appear early in the trial, followed by Cue-coding and Action-coding states. To test this hypothesis, all within-trial onset times of the classified coding states were pooled across all sessions. Trial durations were “warped” so that taste delivery occurred at time 0 and decision occurred at time 1; this way timings could be compared across different trials in common time units. The distributions of coding states’ onset times in these common time units are plotted in **Figure 2a**. The distributions are clearly separated in time and arranged in a sequence where Quality-coding states appear before Cue-coding states, which, in turn, appear before Action-coding states. Mean onset times for Quality-coding, Cue-coding, and Action-coding states were 0.14, 0.41, and 0.67 (common time units), with inferred peaks occurring at 0.07, 0.38, and 0.87, respectively. This coding progression emerges after pooling all trials across all sessions; coding states (specifically, Quality-coding, Cue-coding, or Action-coding) were found in 1, 028 out of 3, 160 total trials, and occurrences of multiple coding states within the same trial were rare, likely due to small neural ensembles per session. However, the ordering of within-trial coding state sequences was highly reliable: out of 362 trials containing at least two different types of coding states, 313 (86%) had the proper ordering (i.e., Quality → Cue → Action). Thus, although the phenomenology is an inferred property of the aggregate pseudo-population of neurons, the results are consistent with sequential encoding in each trial unfolding from sensory information processing to perceptual categorization, and finally to decision-making and action execution.

**Figure 2.**
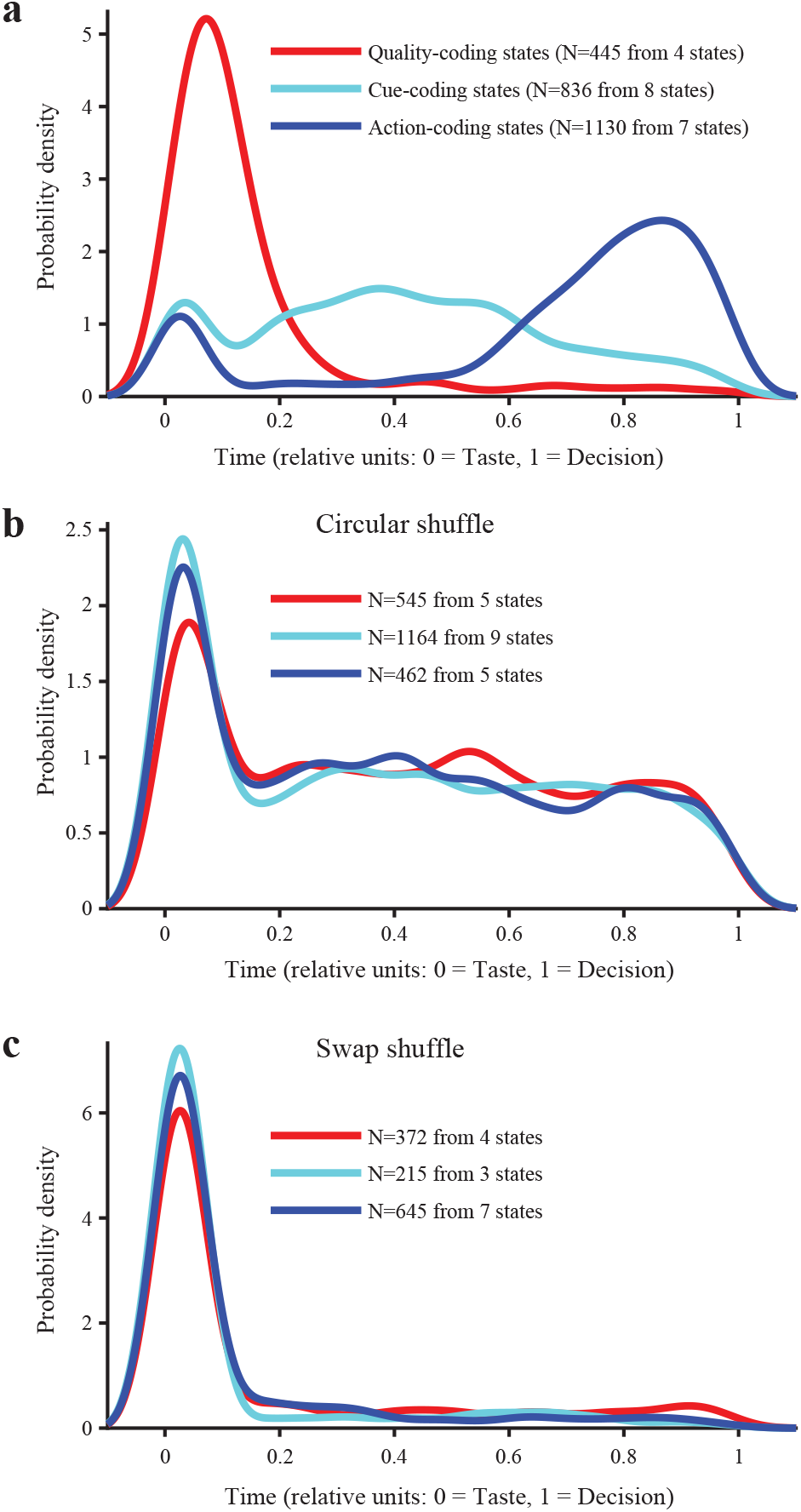
Coding state onset times exhibit an intuitive temporal sequence. **a**: Distribution of onset times of all states classified as either Quality- (red), Cue- (cyan), or Action-coding (blue) for experimental data showing a progression from coding for sensory information to abstract cue information to action information. **b**: Distribution of onset times for spiking data that are shuffled circularly. **c**: Distribution of onset times for spiking data that are swapped across time.

To ensure that coding states were meaningful, and not a by-product of HMM analysis, a series of control analyses were performed. Randomly permuting trial type (i.e., stimulus) labels within correct and incorrect trials and repeating the state coding classification analysis severely reduced the probability of finding coding states in the data, as expected (4 Quality-coding states in unshuffled data vs. an average of 0.3 in shuffled datasets; 15 Decision-coding states in unshuffled data vs. none in shuffled datasets). In addition, two surrogate datasets were generated from raw data by shuffling the spike trains circularly or by swapping segments of simultaneous activity across time (see **Methods**). The circular shuffle preserves individual spike train autocorrelations but disrupts co-activation (hence, pairwise correlations) between neurons; the swap shuffle does the opposite, i.e., it preserves neuron co-activations but disrupts autocorrelations (Maboudi et al., 2018). Both types of shuffling preserve the individual neurons’ firing rates. These shuffled datasets were fitted with new HMMs and the resulting states were re-classified for their coding properties. Overall, we found only a modest reduction in the numbers of different coding states (**Supplementary Table 1**), which may indicate that the neural autocorrelations alone, or the pairwise correlations alone, may contain enough information to identify activity associated with task variables (e.g., if enough information is contained in the firing rates alone). This is possible since (i) the circular shuffling procedure will destroy the HMM states found in the original dataset, but can still produce new states, and (ii) randomly swapping time bins will not disrupt coordinated spiking across neurons. The states produced by shuffling, however, no longer truly reflect task variables, as confirmed by the lack of temporal information in the onset times of the coding states. In particular, the onset time distribution plots constructed for the shuffled datasets showed no clear separation among coding states of different types (**Figure 2b-c**). Thus, even though there was enough information in the shuffled datasets for the model to find coding states, the temporal structure of these states was completely disrupted.

### Modeling metastable decision-making activity in GC

Results from HMM analysis demonstrate that GC relies on metastability to represent variables associated with the performance of a taste-based 2-AC task. To investigate the potential circuit-level mechanisms driving metastable activity in the context of the 2-AC task, we developed a recurrent spiking network model of GC that could replicate the key experimental findings.

Metastability in spiking network models can be achieved by a clustered architecture where synaptic connections among neurons within the same cluster are stronger than those among neurons of different clusters (Deco and Hugues, 2012; Litwin-Kumar and Doiron, 2012; Mazzucato et al., 2015; Rostami et al., 2020). Our network contained both excitatory (E) and inhibitory (I) neurons, each organized into 14 uniformly-sized clusters with stronger synaptic connections than among neurons of different clusters. Each excitatory cluster was paired with a partner inhibitory cluster via stronger synaptic connections than among neurons of non-partner E-I pairs (Rostami et al., 2020) (see **Figure 3a, Supplementary Figure 2a**, and **Methods** for details).

**Figure 3.**
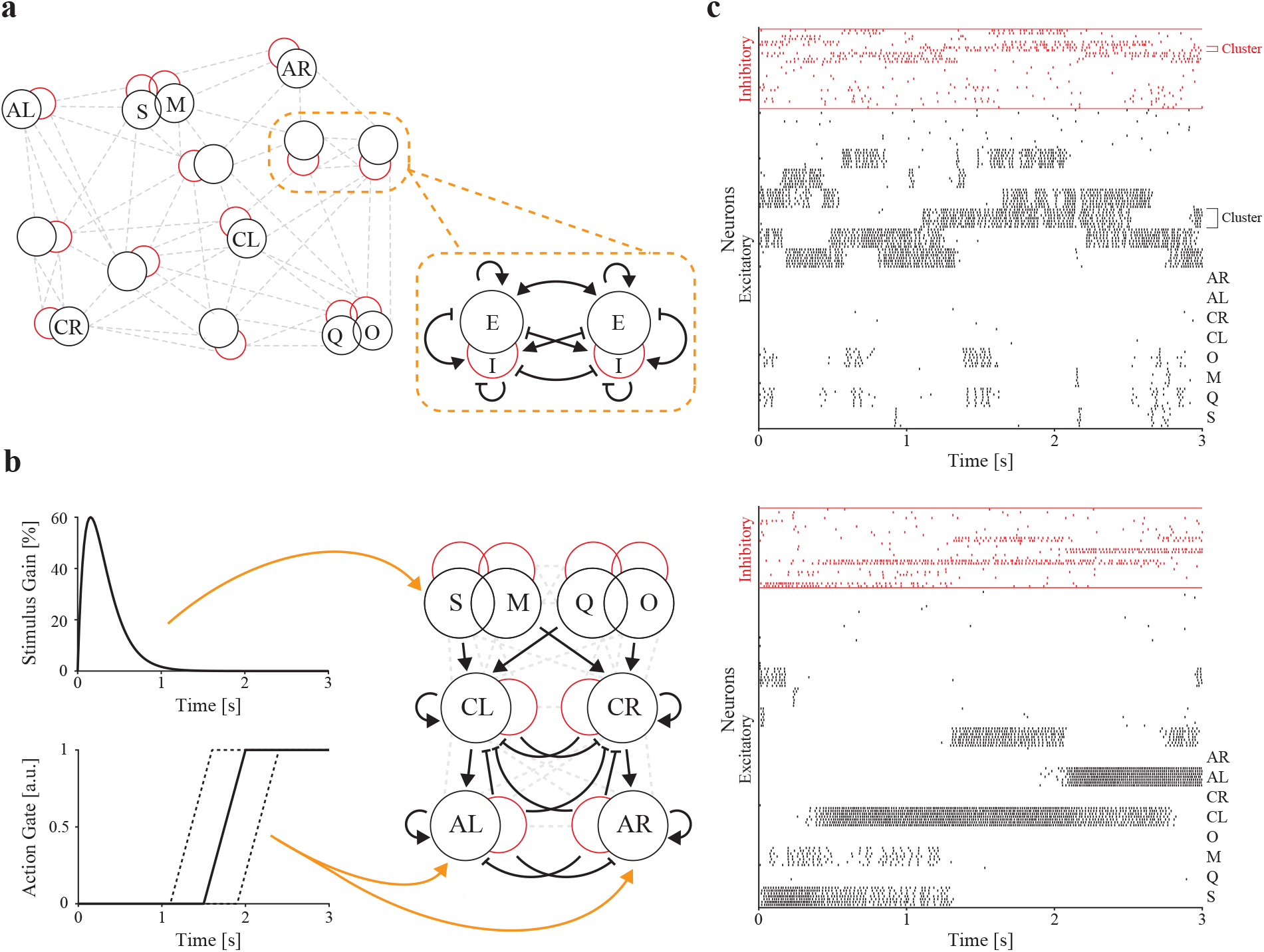
Clustered spiking network model for decision-making activity. **a**: General structure of the network. Neurons in the network are organized into fourteen excitatory (black) and partner inhibitory (red) clusters and connected in a highly recurrent manner. The zoomed inset indicates the generic structure of all of the connections that exist between any two excitatory clusters and their inhibitory cluster partners (pointed arrowheads are excitatory connections, flat arrowheads are inhibitory connections). **b**: Details of synaptic connections between taste, cue and action clusters, with inputs simulating the sensory stimulus (**left, top**) and the gating GO signal (**left, bottom**). **c**: Raster plot of metastable network activity in the absence of a stimulus (**top**) and after stimulation (**bottom**). Neurons are grouped by clusters (black: excitatory, red: inhibitory). S: sucrose, M: maltose, Q: quinine, O: sucrose octaacetate, CL: cue left, CR: cue right, AL: action left, and AR: action right.

The model was built to encode the crucial variables associated with the decision-making task, namely stimulus (taste), cued direction (cue), and selected direction (action). This was achieved by wiring the network such that specific properties could be assigned to 8 randomly selected excitatory clusters, coding respectively for: sucrose, quinine, maltose, sucrose octaacetate, cue left, cue right, action left, and action right. Taste-selective clusters were characterized by having half of the neurons receive additional external input when either sucrose, quinine, maltose, or sucrose octaacetate were presented. The increased input varied in time as an alpha function (difference of two exponentials) that began at the start of the trial (**Figure 3b, top left**).

To replicate the experimental finding of coding states associated with a particular taste quality (i.e., sweet or bitter), clusters of neurons coding for sucrose and maltose were more strongly connected than clusters coding for, e.g., sucrose and quinine. Similarly, clusters coding for quinine and octaacetate were more strongly connected than clusters coding for, e.g., quinine and maltose. The same structure was imposed on the corresponding inhibitory partners. This structure was obtained by randomly selecting one quarter of the neurons in each partner E-I pair and connecting them with strong synaptic weights *J*_++_*J*_*αβ*_, with *J*_*αβ*_ being a common reference synaptic weight from neurons in cluster *β* to neurons in cluster *α* (*α, β* ∈ {E, I}). Neurons in the same taste cluster, but outside this more strongly connected subpopulation, had weaker synaptic connections (mean *J*_+_*J*_*αβ*_, with *J*_++_ *> J*_+_ *>* 1; see **Methods, Supplementary Figure 2b**, and **Supplementary Table 3** for model parameters). This structure mimics overlapping populations of neurons with similar response properties, and reflects the fact that many neurons in the GC (Jezzini et al., 2013) and elsewhere in the cortex (Curti et al., 2004) respond to multiple stimuli.

The 2 cue-selective clusters received stronger input from the appropriate pair of taste-selective clusters (i.e., connections from sucrose and quinine clusters to the “cue left” cluster and from maltose and sucrose octaacetate clusters to the “cue right” cluster were stronger compared to the cross-connections; see **Figure 3b** and **Supplementary Figure 3** for additional details); similarly the 2 action-selective clusters received stronger input from the appropriate cue-selective cluster (i.e., stronger connections from the “cue left” cluster to the “action left” cluster, and from the “cue right” cluster to the “action right” cluster). Cue- and action-selective clusters also had stronger intracluster connections than other clusters (**Supplementary Table 3**) to promote more “persistent” activity intended to reflect stable mental constructs.

To represent the approach of the lateral spouts in the behavior task, which prevents mice from acting on a decision until the end of the delay period, we incorporated a preparatory action gating signal in the model (**Figure 3b, bottom left**). The gate prevents the action-selective clusters from receiving input until a randomly chosen point in time in each trial, representing availability of the spouts. The gating mechanism is implemented as a multiplicative modifier of input that ramps linearly from 0 to 1 over 0.5 s, beginning at a random time (uniformly distributed over [1.1 s, 1.9 s]).

Altogether, the anatomy and topology of the model network gave rise to a random-looking, metastable dynamics in the absence of a stimulus (**Figure 3c, top**), and to a cascade of processes—from taste to cue to action—in the presence of a stimulus (Miller and Katz, 2010), due to preferential connections embedded within the highly recurrent network model (**Figure 3c, bottom**). The network typically responded to stimuli with high firing rates initially in the stimulus-responsive cluster and its overlapping partner, then in a cue cluster, and then in an action cluster (**Figure 3c, bottom**).

To infer the model’s choice on each trial, we computed firing rates for action clusters in 50 ms bins and considered a cluster to be “on” when it fired at ≥ 40% of its maximum firing rate across the whole session. Behavioral scoring of trials was then evaluated following this simple rule: if the correct action cluster came on between 0.5 s and 3.0 s after stimulus onset and the incorrect action cluster did not, the trial was scored as correct; if the incorrect action cluster came on during the same window and the correct action cluster did not, the trial was scored as incorrect; if neither action cluster or both action clusters came on during this window, the decision was assigned randomly.

To test the model, we randomly generated 10 different synaptic weight matrices, which may be thought of as 10 independent network instances. For each instance of the network, we ran 100 trial simulations (25 simulations per tastant); in each simulation, a stimulus was applied and the network was allowed to evolve over 3 seconds. Over all 10 realizations, the network had an average accuracy rate of 80.1%, which is comparable to the experimental performance of mice performing the real task (82.8%; Vincis et al., 2020). To assess whether the network model could reproduce the metastable dynamics observed experimentally, we randomly selected one neuron from each excitatory cluster to form a 14-neuron sample. The spike trains of these sampled 14-neuron ensembles were fitted with an HMM over the 100 trials (**Figure 4a**). Metastable states were decoded on a trial-by-trial basis as was done for the experimental data, and state coding properties were classified according to the same definitions used in **Figure 1b-d** (see **Figure 4c-e**).

**Figure 4.**
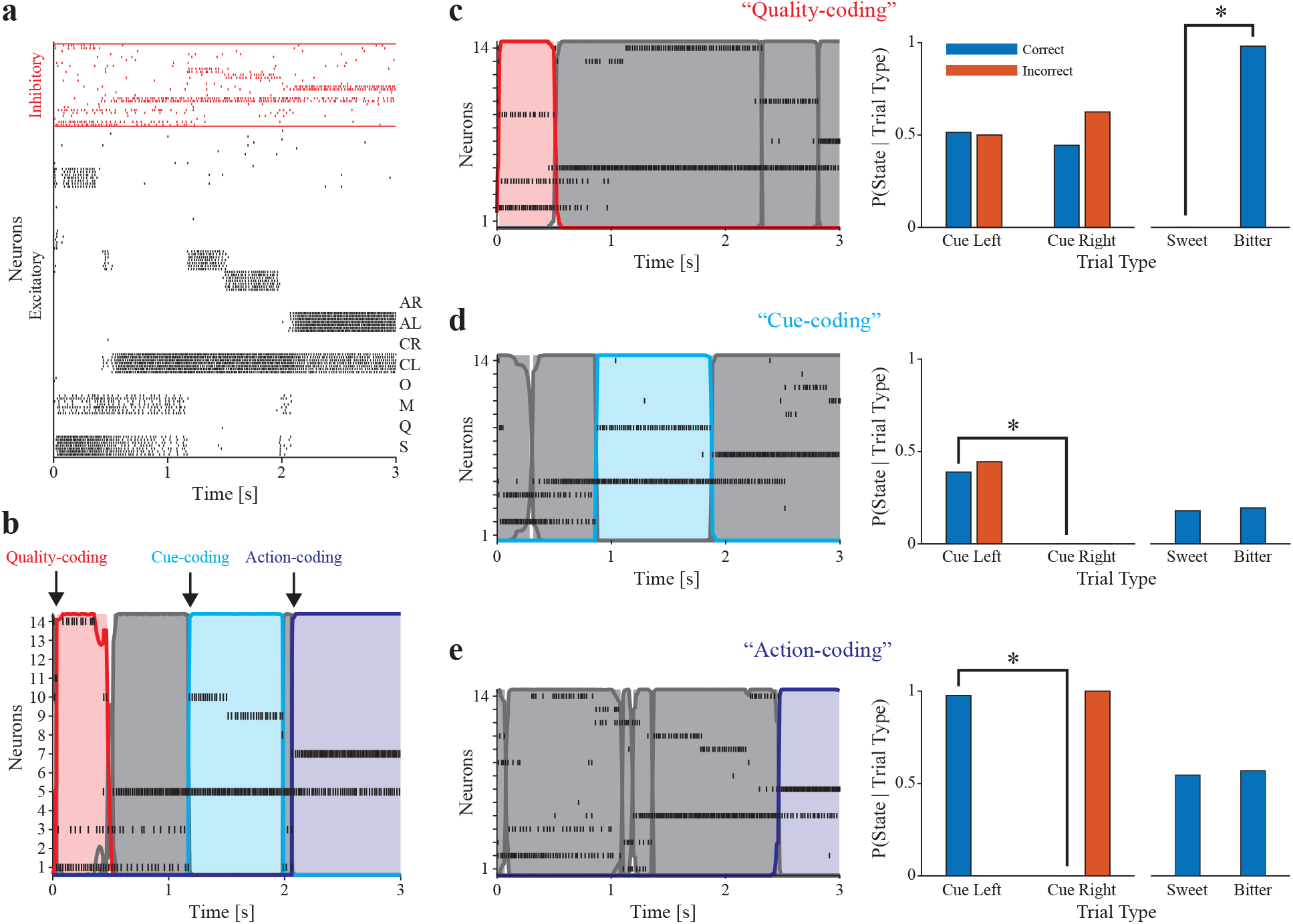
HMM analysis of network-simulated data. **a**: Example network activity showing sucrose-triggered response. **b**: Raster plot with overlaid HMM-decoded hidden states corresponding to the trial shown in **a**. Arrows indicate onsets of different coding states. **c-e (left)**: Raster plots and HMM-decoded metastable states for simulated spiking. **c-e (right)**: Classification of hidden states. Quality-coding (**b**, red), Cue-coding (**c**, cyan), and Action-coding (**d**, blue) states. * indicates significant difference by Chi-squared test (*p* < 0.05, corrected for the number of decoded states in the session).

Analysis of state onset times (**Figure 5a**) revealed that these distributions corresponded quite well with those observed in the experimental data. Most notably, the model captured the temporal sequence of distribution peaks observed in the data, showing a progression through time from taste-coding to abstract cue-coding and finally to action-coding. The mean onset times for Quality-coding, Cue-coding, and Action-coding states were 0.04, 0.49, and 0.74 common time units, with inferred peaks occurring at 0.03, 0.40, and 0.82, respectively. Similarly to the experimental data, no coding states with any meaningful temporal dynamics were found when shuffling the simulated data (circular or swap shuffle, see **Methods** and **Figure 5b-c**). This finding confirms that coding the temporal structure of the task via a sequence of metastable states requires a particular network structure, with our model providing a specific link between network structure and function.

**Figure 5.**
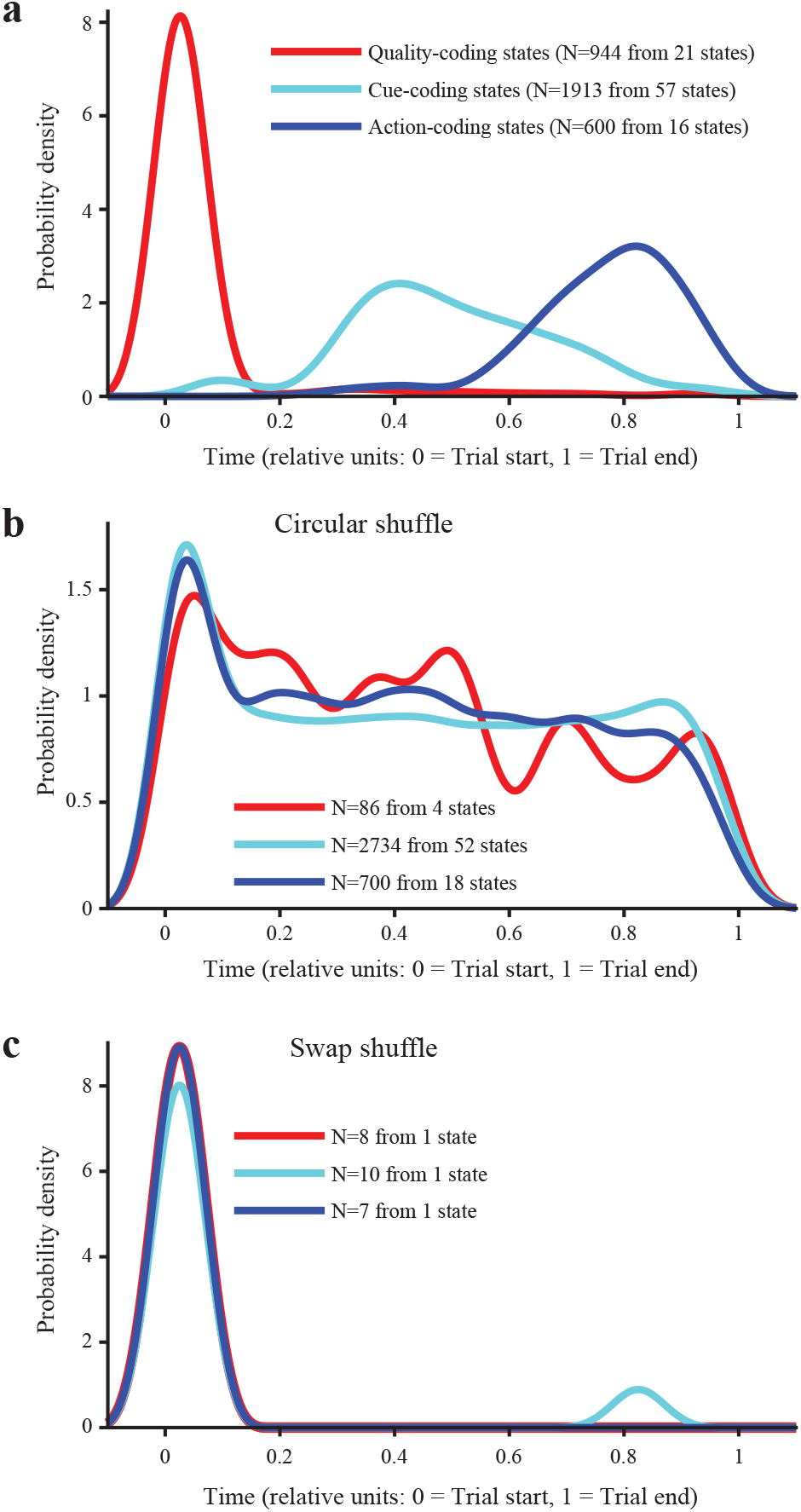
Coding state onset times in the simulated data qualitatively reflect those in experimental data. **a**: Distribution of onset times of all states classified as either Quality- (red), Cue- (cyan), or Action-coding (blue) for simulated data showing a progression from coding for sensory information to abstract cue information to action information. **b**: Distribution of onset times for spiking data that are shuffled circularly. **c**: Distribution of onset times for spiking data that are swapped across time.

### Relationship between GC dynamics and behavioral performance

To further probe the relationship between network structure and function, we subjected our model network to a series of external perturbations, inspired by the optogenetic silencing experiments carried out by Vincis et al. (2020). Those experiments showed that task performance is unaffected by silencing GC during the “sampling period” (from 0 to 0.5 s post-taste event) but impaired by silencing GC during the “delay period” (from 0.5 to 3 s post-taste event). We modeled optogenetic silencing as a simple square pulse input on top of the baseline external current and with strength (height) measured as percent increase in the baseline. The silencing input targeted all inhibitory neurons in the network. Representative spike rasters for network activity with simulated silencing during the sampling and delay periods are shown in **Figure 6a**. We ran 100 simulated trials for each of our 10 networks under both silencing conditions and measured task accuracy as we did before under control conditions. **Supplementary Figure 5a** shows the results of silencing on network performance. Silencing during the delay period induced a more dramatic drop in performance than silencing during the sampling period (*p* < 0.0001; Bonferroni-corrected post-hoc after significant 1-way within-subjects ANOVA); however, silencing during the sampling period also reduced the performance (*p* = 0.0013), contrary to our previous experimental findings (Vincis et al., 2020).

**Figure 6.**
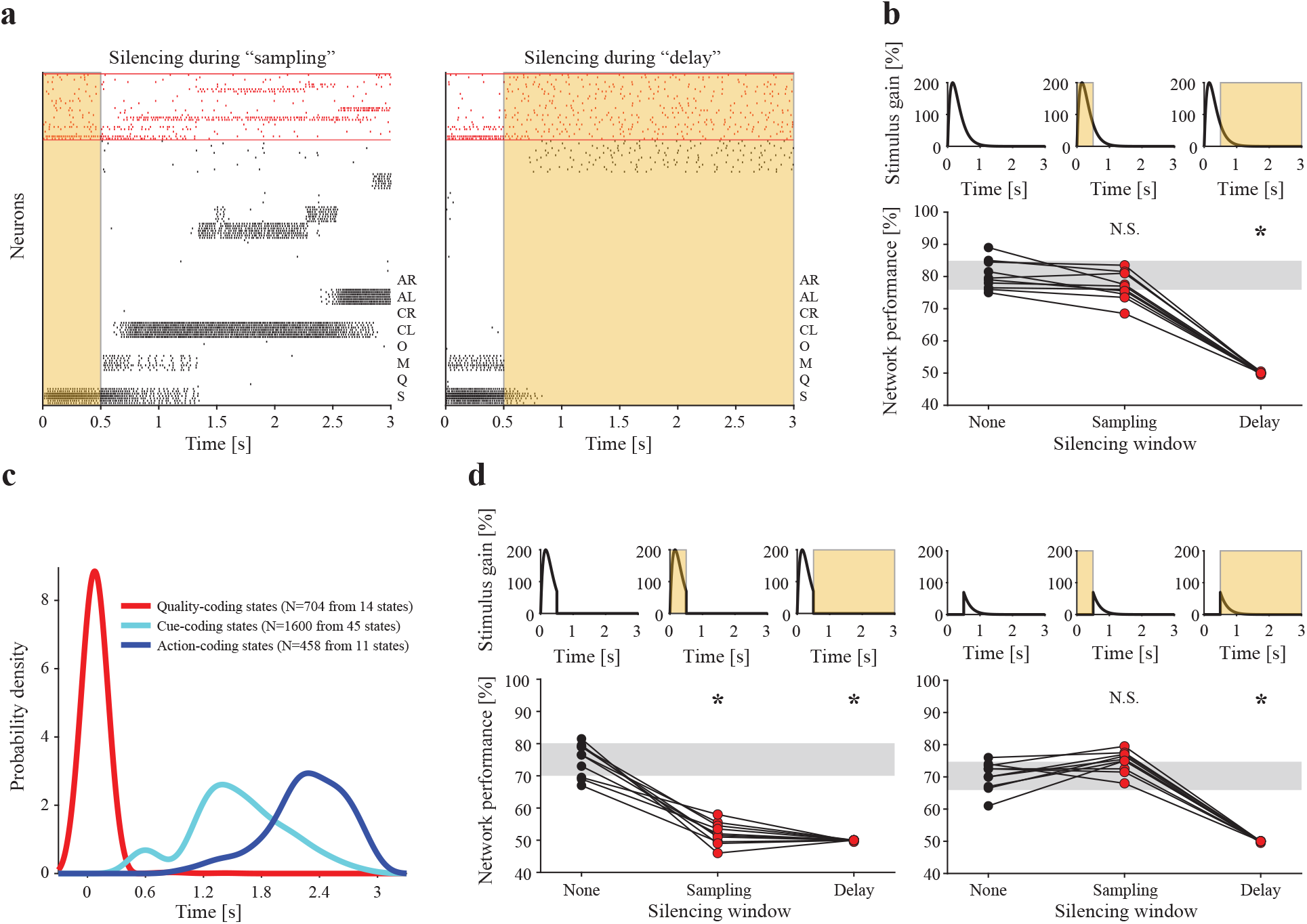
Effect of simulated optogenetic silencing on model performance. **a**: Rasters showing representative network activity when silencing is applied during “sampling” (0 to 0.5 s) and during “delay” (0.5 to 3 s). Shaded yellow regions indicate that silencing (100% increase in external baseline current for all inhibitory neurons) is on. **b**: Stimulus time courses and periods of silencing for 10 networks receiving full stimulus input (**top**) and their corresponding distributions of task accuracies (**bottom**). **c**: Distributions of onset times of coding states for 10 networks with stimulus input with gain of 200% (compare to **Figure 5a**, wherein the stimulus gain is 60%). **d**: Stimulus time courses and periods of silencing for 10 networks receiving partial stimulus input (**top**) and their corresponding distributions of task accuracies (**bottom**). Partial stimulus input was divided into Head only (**left panel**) and Tail only (**right panel**). Grey bars represent means 1 standard deviation for the corresponding None (baseline) condition; N.S. represents no significant difference with respect to the corresponding None; * represents significant difference with respect to the corresponding None (*p* < 0.05, Bonferroni-corrected post-hoc for 2-way within-subjects ANOVA with factors Stimulus and Silencing).

These results depend on the specific parameters used to model the stimulus and the effect of silencing, parameters on which there is little or no experimental information. We therefore adjusted the taste stimulus gain and time constant to capture both the effects of silencing and our HMM results under control conditions. Increasing the stimulus decay time constant from 160 ms to 705 ms yielded performance results that matched the silencing experiments (**Supplementary Figure 5c**), however, repeating the HMM analysis with this new stimulus revealed disrupted coding state dynamics (**Supplementary Figure 5d**). Instead, leaving the decay time constant at 160 ms and increasing the stimulus gain from 60% to 200% captured both the silencing experiments (**Figure 6b**) and the temporal progression of the coding states (**Figure 6c**; compare with **Figure 5c**). Thus, with appropriate stimulus parameters, the model can explain the silencing effects as well as the metastable dynamics of coding states found in the HMM analysis of the data.

With the effect of silencing on behavior now captured by the model, we investigated the mechanism by which silencing during the sampling period fails to impair task performance. There could be two (not mutually exclusive) mechanisms underlying the ineffectiveness of silencing during sampling: (1) the silencing input cannot overcome the simultaneous, strong stimulus input during the first half second; (2) the residual stimulus input occurring after the first half second provides sufficient information to drive network performance. We tested these hypotheses by decomposing the stimulus into two segments: the “head” that occurs during the first half second and the “tail” that occurs after the first half second. We then conducted additional simulations to measure network accuracy under control and under both silencing conditions for each component of the stimulus (**Figure 6d**). We compared performance values using a 2-factor within-subjects ANOVA with two 3-level categorical factors: Taste Stimulus (levels: Full, Head, and Tail) and Silencing (levels: None, Sampling, and Delay). We found a significant Stimulus-Silencing interaction and carried out post-hoc comparisons with a Bonferroni correction. Although each stimulus component was necessary for normal performance when no silencing was present (Full/None vs. Head/None, *p* = 0.0026; Full/None vs. Tail/None, *p* < 0.0001), the tail of the stimulus was both necessary and sufficient for normal performance in the presence of silencing during the sampling period (Full/Sampling vs. Head/Sampling, *p* < 0.0001; Full/Sampling vs. Tail/Sampling, *p* = 0.6572). This supports the latter of the two explanations described above—i.e., that residual stimulus input occurring after the first half second is sufficient for accurate performance. This also suggests that the dynamics of lingering stimulus information in GC can rescue the silencing-induced performance deficits after the sampling period.

To further explore this time-variable effect of silencing with a higher resolution, we took a 250 ms-wide silencing square pulse and slid its center along the 3 s trial window in 50 ms increments, computing the 10-network, 100-trial average performance at each point. Two silencing strengths were used: “Weak” (25%) and “Strong” (100%). This revealed 3 temporal windows of interest: i) sampling onset, when both weak and strong silencing have no effect on performance; ii) sampling offset and cue cluster onset, when both weak and strong silencing impair performance; and iii) mid-trial, when strong silencing impairs performance but weak silencing does not (**Supplementary Figure 6**). We investigated the impact of silencing in these three windows on network activity by taking a representative network and comparing its control trials (i.e., those without silencing) to silenced trials. We used the same initial conditions in all cases and 6 different combinations of silencing perturbation: Weak or Strong at the beginning of the sampling (“Beginning”: 0 to 250 ms), Weak or Strong at cue onset (“Cue onset”: 250 ms window centered on the onset time of the cue cluster in the corresponding control trial—these onset times have mean ∼ 0.6 s), and Weak or Strong during the middle trial period (“Middle”: 1, 375 to 1, 625 ms). We fit an HMM to the 700 total trials of spiking activity generated by this network, and decoded its hidden states in each trial (**Figure 7a**). Effects on metastable states paralleled the effects on network task performance. Visually, the sequences of states under the control condition appear similar to those under silencing conditions that did not impact performance (Weak/Beginning, Strong/Beginning, and Weak/Middle) and different from those under silencing conditions that did impact performance (Weak/Cue onset, Strong/Cue onset, and Strong/Middle), wherein state transitions are much more frequent and state sequences appear muddled. This phenomenon is confirmed by computing an average pairwise state matching rate relative to control conditions (see **Methods**) as a measure of sequence similarity to control sequences over time (**Figure 7b**). Classifying the hidden states further reveals a significant decrease in the fraction of trials containing a correct Action-coding state under the three silencing conditions that impair performance (*p* < 0.05 for Marascuilo post-hoc tests vs. control after significant Chi-square test across conditions; **Figure 7c**).

**Figure 7.**
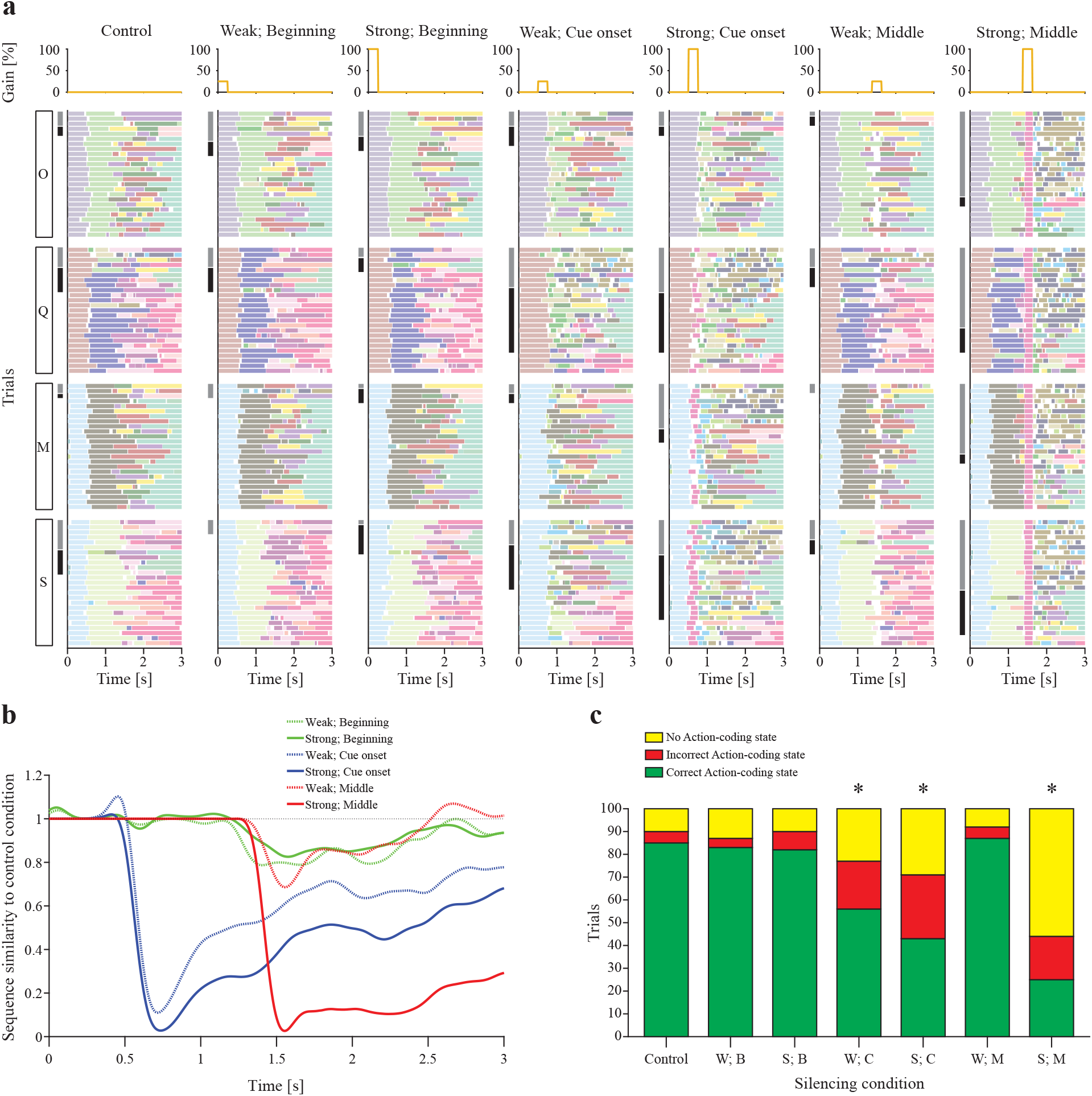
Effect of simulated optogenetic silencing on network activity. **a**: HMM-decoded states for a network subjected to various simulated optogenetic perturbation conditions. Silencing time courses for each condition are shown at the top of each column; decoded states over all 100 trials for each condition are shown below. Trials are ordered in stimulus blocks from bottom to top (S: sucrose, M: maltose, Q: quinine, O: octaacetate) and are ordered in outcome blocks within each stimulus block from bottom to top (correct, incorrect, omitted; incorrect and omitted trials are indicated by black and grey shading, respectively, on the left-hand side). **b**: Average state sequence similarities over time. For each silencing condition in **a**, the corresponding curve represents how similar the state sequence is (on average) to the state sequence obtained under control (i.e., under no silencing). Each raw similarity score (ranging from 0 to 1) was normalized by the score obtained by comparing the control condition with itself (see **Methods** for details, Eq. 5). **c**: Numbers of trials out of 100 for each condition (W: Weak, S: Strong, B: Beginning, C: Cue onset, M: Middle) that contain correct, incorrect, or no Action-coding states, based on a coding state classification of the states from **a**. Action-coding states are considered correct or incorrect in each trial based on whether their direction preference (in correct trials) matches that trial’s correct direction. * indicates significant difference vs. control in fraction of trials with correct Action-coding states (Chi-squared test across conditions, followed by pairwise Marascuilo post-hoc with *α* = 0.05).

## Discussion

In this study, we demonstrate that neural activity associated with taste-based decisions is metastable and temporally organized to sequentially encode information related to sensory stimuli, decision variables, and upcoming actions. We further provide a computational model that accounts for metastable dynamics in GC and its link with behavior. It further shows the relationship between silencing, metastable dynamics, and network performance. This model allows us to explain why silencing GC during sensory stimulation has a smaller impact on performance compared to perturbation during the deliberation period. These results significantly advance our understanding of GC and its role in decision-making, and confirm metastability as a dynamic regime relevant for GC’s function.

### Temporal dynamics of GC activity and functional roles of metastability in GC

GC neurons produce time-varying responses following gustatory stimulation (Katz et al., 2001; Katz et al., 2002; Bouaichi and Vincis, 2020). Firing rates transition through three successive epochs, each having a different onset and encoding a distinct aspect of taste. The earliest epoch lasts for ∼ 250 ms and encodes the contact of taste with the tongue. The second epoch (from ∼ 250 ms to ∼ 1 s) represents chemosensory information—that is, taste quality (i.e., bitter, sweet, salty, sour, and savory). Finally, the third epoch (after ∼ 1 s) encodes the palatability of the stimulus and correlates with ingestive or aversive behaviors. This sequence is believed to result from recurrent interactions among GC, thalamic, and limbic areas such as the basolateral nucleus of the amygdala (Piette et al., 2012; Samuelsen et al., 2012; Samuelsen et al., 2013; Livneh et al., 2017; Lin et al., 2021). These dynamics were initially uncovered through a careful analysis of single neuron firing rate modulations averaged across trials (Katz et al., 2001). Additional studies of the ensemble activity from simultaneously recorded neurons further confirmed the existence of these dynamics at the single trial level, revealed trial-by-trial consistency of epochs, and identified epochs as metastable states with rapid onsets (Jones et al., 2007; Mazzucato et al., 2015; Mazzucato et al., 2016).

Recent experiments in rodents engaged in behavioral tasks suggest that additional epochs containing task-dependent variables could exist. For instance, recordings from GC of rodents intentionally licking a spout show preparatory activity that precedes the somatosensory epoch (Stapleton et al., 2007; Dikecligil et al., 2020). Studies using taste-anticipating cues prove the existence in GC of anticipatory firing rate modulations, which encode general and specific expectations (Samuelsen et al., 2012; Gardner and Fontanini, 2014; Livneh et al., 2017). Similarly, experiments in rodents performing a decision-making task show neural activity consistent with sequences of epochs encoding decision-related variables (Fonseca et al., 2018; Vincis et al., 2020). While this more recent work has been relevant in showing that GC does not just multiplex sensory variables, it has largely relied on single neuron analyses and across-trial averages.

The work presented here is the first that shows that variables associated with reward-driven decision-making are encoded by GC as metastable states. Our HMM analysis of GC ensemble activity during the four-taste, perceptual decision-making task has demonstrated that GC activity traverses sequences of metastable states encoding relevant task variables. These results are in line with prior evidence from higher order areas (Durstewitz et al., 2010; Ponce-Alvarez et al., 2012; Stokes et al., 2013; Rich and Wallis, 2016; Benozzo et al., 2021) and are the first to show such a phenomenon in a sensory area. Our analysis further shows that, in the rodent GC, metastable states undergo a temporal progression with taste-coding states occurring first, followed by cue-coding states, and finally by action-coding states. This progression is consistent with a model in which GC first encodes sensory information (e.g., sweet vs. bitter), then decisional variables (e.g., predictive value of taste or preparatory activity associated with actions), and then actions (e.g., lick left vs. lick right). Importantly, coding state transitions occur at variable times in different trials, allowing for pinpointing of the internal occurrence time of sensory perception or decision. This also allows for disentangling of the neural activity related to internal processes from the neural activity immediately following external triggers.

### Perturbing GC dynamics

GC dynamics and their behavioral significance have been extensively tested via temporally precise optogenetic perturbations. Studies in rats receiving tastants through intraoral cannulae demonstrated that the effects of perturbations on gaping depend on the specific timing of silencing (Mukherjee et al., 2019). The differential impact of perturbations during specific temporal epochs was also present in mice performing a four-taste, perceptual decision-making task (Vincis et al., 2020). Optogenetic activation of parvalbumin-positive interneurons—an established method for silencing cortical activity (Guo et al., 2014)—was effective in reducing behavioral accuracy only if performed during the delay period, when cue-related, preparatory activity occurs, and not during the period of taste sampling. While these results were interpreted as evidence that GC’s cognitive signals play a role in guiding taste-based decision-making, they also raised questions about the behavioral role of GC taste-coding states. In this work, we capture those results in our spiking network model and provide a possible explanation. Our results suggest that the ineffectiveness of silencing during the sampling period is best explained by residual stimulus input sufficing to drive the network after silencing, although competition between simultaneous stimulus and silencing inputs may also play a role.

Our additional silencing perturbation simulations, conducted at higher temporal resolution than explored experimentally, revealed precise points of susceptibility and resilience to silencing in the model, both in terms of task accuracy and metastable state sequences (**Supplementary Figure 6**). Model activity, metastable dynamics, and performance were robust against silencing when it occurred at the beginning of a trial, were disrupted by silencing when it was applied at the onset time of a cue cluster, and—depending on silencing strength—were conditionally impaired by silencing when it occurred in the middle of a trial (**Figure 7**). Furthermore, metastable dynamics and their ability to sequentially encode stimulus and task variables were tightly linked to network performance. The model therefore provides experimentally testable predictions for optogenetic silencing of neural decision-making activity with various intensities and windows of application.

### Modeling GC: Metastability and decision-making

The observation of reliable and reproducible metastable dynamics in GC has inspired modeling efforts aimed at explaining the network mechanisms underlying this phenomenon and its functional significance. Miller and Katz (2010) showed how a spiking metastable network can account for state sequences observed during consummatory behavior (associated with ingestion or rejection of tastants); later, Mazzucato et al. (2015) demonstrated the existence of metastable dynamics in the absence of overt stimulation, and built a spiking network model of GC able to spontaneously produce metastable dynamics. This network has formed the basis of subsequent models that have captured a wealth of data pertaining to neural dynamics in GC (Mazzucato et al., 2016; Mazzucato et al., 2019). The model described here represents another significant advancement, in that the model can further process stimuli based on the abstract decision they instruct. The four tastants instructing decisions are categorized according to their taste quality first and their predictive value after, reproducing the temporal sequence of metastable states representing taste quality, predictive value of the taste, and action. The model also produces decisions with a similar performance to our experimental subjects.

In order to reproduce the experimental results observed in mice performing the four-taste, 2-AC task, the network had to include the following features: i) partial overlapping of clusters representing stimuli that share the same taste quality; ii) biased intercluster connections creating a flow of information from sensory clusters to cue clusters to action clusters; iii) an action gate signal priming action. These ingredients were absent in previous models and suggest the existence of possible biological counterparts in GC. Partially-overlapping clusters take into account that GC neurons may respond to multiple tastes (Jezzini et al., 2013) and may be the basis of stimulus generalization based on predominant taste quality. Biased intercluster connectivity may be the result of synaptic plasticity leading to the formation of neuron ensembles producing preparatory and action-related activity. The action gate was inspired by the presence of the lateral spouts moving toward the mouse, hence providing an implicit, ramping “GO” signal that may correspond either to an anticipatory input from areas like the amygdala, or to a disinhibitory signal triggered by the cue. This gate is also similar in spirit and implementation to the “urgency-gating model” described by Cisek et al. (2009). Additional experiments are needed to test these predicted mechanisms.

The model also explains the effects of optogenetic perturbations and makes further predictions based on the strength of the perturbation and the period in which it is applied, as discussed in the previous section.

In summary, the present work shows that metastable dynamics occurs in GC during decision-making and provides a computational model explaining the genesis of metastability and its role in taste-based decisions. The ability of our model to reproduce the experimental results, together with our past body of work (Mazzucato et al., 2015; Mazzucato et al., 2016; Mazzucato et al., 2019), suggest that the clustered inhibitory-excitatory network model may truly underlie the anatomical and functional organization of the rodent GC.

## Methods

### Summary of behavioral procedure and electrophysiology

Methods for behavior and electrophysiology are available in Vincis et al. (2020), which presents the dataset analyzed in this study. Briefly, mice were unilaterally implanted with 8 tetrodes in the left gustatory cortex for chronic electrophysiological recording. After recovering from surgery, mice were water-restricted and trained to perform a four-taste, perceptual decision-making task to obtain water rewards. For the task, head-fixed mice sample a tastant—either sucrose, quinine, maltose, or sucrose octaacetate—from a central spout. The sampling period (time between first and last lick to the central spout) was ∼ 0.5 s. After a delay period (∼ 2 s) two lateral spouts advance in front of the animal’s mouth. To receive the water reward, the mouse must lick the left spout if the tastant was sucrose or quinine and must lick the right one if the tastant was maltose or sucrose octaacetate. Testing commenced once mice achieved a criterion performance of at least 75% for 3 consecutive days. Electrophysiological recordings were obtained while mice performed the task.

### Raw data and preprocessing

Raw data consisted of spike trains from simultaneously recorded neurons over 21 recording sessions involving 5 mice. Within each recording session, trained mice performed an average of 190 trials (range: 75 − 292) of the behavioral task at an average accuracy of 82.8% (range: 71.0% − 93.3%). Only spikes within 100 ms before the taste event (first lick to central spout) to 100 ms after the decision event (first lick to lateral spout) were considered for each trial (we refer to these 100 ms segments before and after events as “padding”). These windows of analysis for each trial were concatenated and, prior to model fitting, neurons whose firing rates averaged over this total window were *<* 2 Hz were excluded from analysis. Sessions with < 3 simultaneously recorded neurons were then excluded from analysis. This resulted in 16 recording sessions involving 4 mice, with an average of 5.4 neurons per session (median: 5.5; range: 3 − 9).

### Model fitting and hidden state decoding

Hidden Markov Models (HMMs) were fit to spiking data from Vincis et al. (2020) as described in Mazzucato et al. (2019). Briefly, spike sequences in a single session were binned into 2 ms bins, and each bin was assigned a symbol from 1 to *N* + 1, where *N* is the number of neurons and symbol s means that only the sth neuron fired in the current bin (multiple firings were rare and were replaced with a random selection of one of the firing neurons). The HMM is completely determined by the number of hidden states, *M*, the transition probability matrix, *T*, and the emission probability matrix, *E*. The entry *T*_*ij*_ of the *M* × *M* transition probability matrix is the probability of making a transition from state *i* to state *j*; the entry *E*_*is*_ of the *M* × (*M* + 1) emission probability matrix is the probability of producing symbol *s* while in state *i*. This model can be considered an approximation to a Poisson-HMM in the case of very short time bins, with the exception that firing of multiple neurons in the same bin is not permitted (see Mazzucato et al., 2015 for details). The HMMs were fitted to the data by maximum likelihood, using the standard Baum-Welch expectation-maximization algorithm (Rabiner, 1989). For each dataset, we fitted HMMs with values of *M* ranging from 2 to 50 and then chose the model with the lowest BIC score as the best model (see below). The maximal number of hidden states selected in the best model never exceeded 29. For each value of *M*, 10 different random initial conditions were used for *T* and *E*. For initial values of *T*, off-diagonal entries were close to zero and diagonal terms were close to 1. For initial values of *E*, entries were drawn independently from a uniform distribution between 0 and 1.

From each initial condition, the Baum-Welch algorithm was performed for 50 iterations (or until convergence at < 10^*−*10^ tolerance on the log-likelihood, transition, and emission probabilities). Out of the 10 models, the model with the highest log-likelihood was selected as the best model for that given value of *M*. The best models for different *M* values were then compared using the Bayesian Information Criterion (BIC) to obtain the optimal model with the optimal number of states (Ponce-Alvarez et al., 2012; Mazzucato et al., 2019; Benozzo et al., 2021). Specifically, the optimal model was the one that minimized the BIC score: −2*LL*+[*M* (*M* −1)+*MN*]ln*B*, where *LL* is the log-likelihood of the model given the data, *M* is the number of hidden states, *N* is the number of neurons, and *B* is the total number of observations in each session (i.e., the total number of time bins across all trials).

After fitting the models, we computed the posterior probabilities of each hidden state given the model and the data in each bin in single trials. A state admissibility criterion was then imposed: a state was considered detected with enough confidence only if its posterior probability was ≥ 0.8 for ≥ 50 consecutive ms (Mazzucato et al., 2019; see also Seidemann et al., 1996; Jones et al., 2007; Miller and Katz, 2010; Bollimunta et al., 2012; Ponce-Alvarez et al., 2012; Mazzucato et al., 2015; Benozzo et al., 2021 for similar methods).

### Classification of decoded states

For each session, all trials were associated with two labels: a stimulus label (sucrose, quinine, maltose, or sucrose octaacetate) and an outcome label (correct or incorrect). This allowed us to compute occurrence probabilities of states given the trial labels, *P* (*S* | Stimulus, Outcome), as the fraction of trials with the given labels in which state *S* was detected at least once. Each decoded state’s coding properties were determined by first looking only at correct trials and comparing the state occurrence probability *P* (*S* | Stimulus, Correct) across all 4 stimulus labels through a Chi-squared test. If significant, all 6 pairwise post-hoc Marascuilo tests (Prins, 2013) were examined. If there was exactly one stimulus label for which all post-hoc tests involving it were significant, *S* was classified as a Taste ID-Coding state. Otherwise, coding properties were investigated by grouping all trials into cue left vs. cue right trials (sucrose and quinine vs. maltose and sucrose octaacetate trials) and sweet vs. bitter trials (sucrose and maltose vs. quinine and sucrose octaacetate trials). The fraction of cue left, correct trials featuring *S, P* (*S* | Cue left, Correct), was compared to the fraction of cue right, correct trials featuring *S, P* (*S* | Cue right, Correct) through a second Chi-squared test, and the fraction of sweet, correct trials featuring *S, P* (*S* | Sweet, Correct), was compared to the fraction of bitter, correct trials featuring *S, P* (*S* | Bitter, Correct), through a third Chi-squared test (see **Supplementary Figure 1**). In each case, significance level was taken to be 0.05, corrected for the number of decoded states, *N*_d_ (i.e., *p* < 1 − (1 − 0.05)^1*/N*^d is considered significant). Results from these tests were used to classify states as Decision-coding (significant result on second test but not on third test), Quality-coding (significant result on third test but not on second test), Dual-coding (significant results on both tests), and Non-coding (non-significant results on both tests). Decision-coding states were further divided into Cue-coding and Action-coding, as explained below. To control for chance occurrences of coding states, stimulus labels on correct trials were randomly permuted 10 times and, following each permutation, proportions of trials containing each decoded state were re-computed and the same Chi-squared tests were re-run to re-classify the states. We then compared the numbers of coding states found with this shuffling procedure to the number found from unshuffled data.

Decision-coding states were further subclassified by whether their “preferred direction” switched or remained the same between correct and incorrect trials. A coding state’s preferred direction was defined, separately for correct and incorrect trials, as the cued direction that maximized its occurrence frequency (assuming this frequency differed between trials of opposite cued directions): if *P* (*S* | Cue left) > *P* (*S* | Cue right) in correct and incorrect trials, Decision-coding state *S* codes for the left cued direction, and analogously for states coding for the right cued direction. Vice versa, if *P* (*S* | Cue left) > *P* (*S* | Cue right) in correct trials, but *P* (*S* | Cue left) < *P* (*S* | Cue right) in incorrect trials, Decision-coding state *S* codes for the action taken; the same conclusion is reached if *P* (*S* | Cue left) < *P* (*S* | Cue right) in correct trials, but *P* (*S* | Cue left) > *P* (*S* | Cue right) in incorrect trials. Finally, if the state *S* occurred equally often in cue left and cue right trials (either for correct or incorrect trials), we did not subclassify the Decision-coding state.

### Randomization controls

Prior to model fitting, raw data were manipulated to form two additional surrogate data sets that served as additional controls (Maboudi et al., 2018; Benozzo et al., 2021; Recanatesi et al., 2021). For the first control, each spike train was shuffled circularly in each trial of a given session: a random number ∆*t* from the uniform distribution on [0, *t*_trial_], where *t*_trial_ is the time length of the trial (including the padding), was added to each neuron’s spike train in each trial independently such that the portion of the spike train that now exceeded the end of the trial would “wrap back around” to the beginning (i.e., if a trial window starts at 0 and ends at *t*_trial_, and the spike train {*t*_*i*_} is shifted by ∆*t*, the new spike train is {*t*_*j*_} = {*t*_*i*_ + ∆*t*, if *t*_*i*_ + ∆*t* < t_trial_; *t*_*i*_ + ∆*t* − *t*_trial_, otherwise}). This procedure preserves the autocorrelations of individual neurons but alters the co-activation/pairwise correlations between neurons. For the second control, each session had its raw spiking data swap-shuffled: each trial’s neuron-by-time spike raster was segmented into 5 ms bins and then these bins/columns were randomly permuted for each trial independently. This randomization procedure preserves the co-activation/pairwise correlations between neurons but removes the autocorrelations of individual neurons. Both procedures preserve the trial-averaged firing rates of individual neurons.

### Onset time analysis

For each session, lists of decoded states of each classified type (Quality-coding, Cue-coding, and Action-coding) were obtained. For each state in each list, all onset times of the state in each trial were obtained. Onset times were expressed relative to their trial’s taste event (*t* = 0), with any negative times (i.e., states beginning during the left padding period) taken to be 0. To account for the fact that the inter-event interval (IEI) between the taste and decision event varied across trials (average: 2.52 s; range: 2.01 − 6.66 s), time was rescaled (or “warped”) in each trial by dividing onset times by the value of IEI in that trial. As a result, warped onset times were between 0 and 1 (where 0 is the time of the taste event and 1 is the time of the decision event) regardless of trial, neuron, session, and classification of the hidden state, and could be directly compared. From this, we obtained a list of (warped) onset times for coding states of each classified type. These lists were pooled across all sessions to obtain histograms of the onset times for coding states of each classified type (bin size: 0.05 warped time units). From the histograms, probability distributions of the onset times were estimated from linear interpolation on the peaks (100 points between any two adjacent bins). Distributions were then smoothed with a Gaussian filter (400 point width).

### Spiking network model

The core of the model was a clustered excitatory/inhibitory spiking network with 4, 000 excitatory (E) neurons and 994 inhibitory (I) leaky integrate-and-fire (LIF) neurons. Each neuron *i* had subthreshold dynamics that obeyed the differential equation in time *t*:

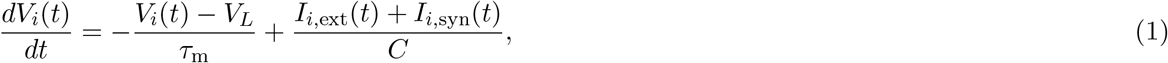

where *V*_*i*_(*t*) is neuron *i*’s membrane potential, *V*_*L*_ its equilibrium/leak potential, *τ*_m_ its membrane time constant, *C* its capacitance, and its total current *I*_*i*_(*t*) is the sum of its external input current *I*_*i*,ext_(*t*) and its synaptic current *I*_*i*,syn_(*t*). The synaptic current to neuron *i* in turn obeyed the differential equation in time *t*:

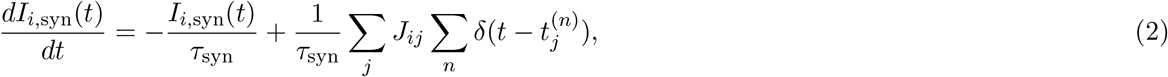

where *τ*_syn_ is the synaptic time constant (different for excitatory and inhibitory presynaptic neurons), *J*_*ij*_ is the synaptic connection strength from presynaptic neuron *j, δ* is the Dirac delta function, and 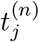 is a presynaptic spike time with *n* indexing over all spikes from a particular presynaptic neuron *j*. This system of differential equations was integrated numerically using the forward Euler algorithm with time step *dt* = 0.05 ms. Whenever *V*_*i*_(*t*) hit a threshold *V*_th_, a spike was emitted and the membrane potential was immediately reset to a value *V*_r_ for a refractory period *τ*_r_ (see **Supplementary Table 3** for parameter values).

The E neurons were arranged into 14 clusters of 250 neurons each, plus 500 neurons not arranged in clusters (the “background population”). The I neurons were arranged into 14 clusters of 71 neurons each. Neurons were randomly connected with probability *P*_*αβ*_, with *α, β* ∈ {E, I}. In general, neurons in the same cluster had stronger synaptic connections than neurons belonging to separate clusters, given by *J*_E++_*J*_EE_ inside E clusters and *J*_I++_*J*_II_ inside I clusters. Each E cluster was paired with a partner I cluster. Synaptic weights between E and I neurons belonging to E-I partner clusters (*J*_I+_*J*_*αβ*_, with *α, β* ∈ {E, I}) were stronger than between E and I neurons in non-partner E-I clusters (*J*_I*−*_*J*_*αβ*_, with *α, β* ∈ {E, I}) (see **Supplementary Figure 2a** and **Supplementary Table 3** for network parameters).

Eight out of the 14 excitatory clusters were randomly selected and assigned “roles” to represent crucial task information: taste clusters (sucrose, quinine, maltose, and sucrose octaacetate), cue clusters (cue left and cue right), and action clusters (action left and action right). For each taste cluster, half of the neurons inside the cluster responded to additional external input when that particular stimulus was presented to the network. All presented stimuli had the functional form 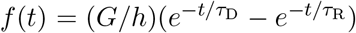, where *G* is the maximum gain, *τ*_D_ and *τ*_R_ are the decay and rise time constants, respectively, of the function 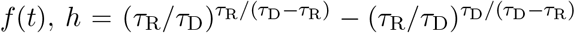 is a normalization constant, and the value of the function determined the percentage increase in external current at time *t* for selective neurons in the appropriate cluster (i.e., 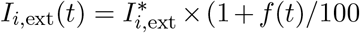) for the constant baseline external current 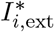).

Unlike generic clusters (i.e., clusters not ascribed a role), only 25% of neurons in a taste cluster were strongly connected (via synapses *J*_E++_*J*_EE_ or *J*_I++_*J*_II_, see above), while the remaining 75% were less strongly connected (*J*_E+_*J*_EE_ or *J*_I+_*J*_II_, respectively for E and I clusters, with *J*_++_ *> J*_+_ *>* 1; see **Supplementary Figure 2b**). The subcluster of taste neurons with strong interconnections was also strongly connected with the subcluster of neurons representing the same taste quality (sweet or bitter), i.e., the sucrose and the maltose subclusters were connected with synapses equal to *J*_E++_*J*_EE_ or *J*_I++_*J*_II_ (for E or I neurons, respectively), and the same was true for the quinine and sucrose octaacetate subclusters. The rest of the connections were as for generic clusters shown in **Supplementary Figure 2a**. As shown in **Supplementary Figure 2b (top)**, it may help to think of this structure as introducing overlaps between taste clusters that share the same taste quality (i.e., sweet or bitter).

Excitatory cue clusters had stronger synaptic connections from the appropriate E taste clusters. For example, 60% of the weights from sucrose and quinine clusters to the cue left cluster and 60% of the weights from maltose and sucrose octaacetate clusters to the cue right cluster were increased by 85% (*J*_CT,E_*J*_E*−*_*J*_EE_, see **Supplementary Figure 3** and **Supplementary Table 3** for parameter values). To facilitate longer lasting representations of cue information in the model, cue clusters had 60% of their excitatory intracluster weights scaled up by 5% (*J*_CC,E_*J*_E++_*J*_EE_), and their synaptic time constants (*τ*_syn,CueE_, *τ*_syn,CueI_) increased (2.3 times greater for excitatory clusters, 1.3 times greater for their inhibitory partners). To discourage simultaneous activation of both cue clusters, 50% of the connections from inhibitory partner clusters of one cue direction to excitatory clusters of the opposite cue direction (*J*_CC,I_*J*_I*−*_*J*_EI_) were increased by 40%.

Excitatory action clusters had potentiated connections from E neurons in cue clusters, with a preference for the appropriate direction: 50% of the weights from, e.g., cue right to action left were multiplied by 2.60 (*J*_AC,Inc_*J*_E*−*_*J*_EE_), while 50% of the weights from, e.g., cue left to action left were multiplied by 2.75 (*J*_AC,Cor_*J*_E*−*_*J*_EE_). We encouraged action clusters, similar to cue clusters, to represent information over longer time scales by scaling up 50% of their intracluster weights by 7% (*J*_AA,E_*J*_E++_*J*_EE_) and increasingtheir synaptic time constants (*τ*_syn,ActE_ and *τ*_syn,ActI_) 3.30 times for excitatory clusters and 3.25 times for their inhibitory partners. As done with cue clusters, we discouraged co-activation of action clusters by increasing the strength of 50% of the connections from inhibitory partner clusters of one action direction to excitatory clusters of the opposite action direction three-fold (*J*_AA,I_*J*_I*−*_*J*_EI_). Finally, to promote a transfer of information from cue to action via negative feedback, we increased the strength of 50% of the connections from inhibitory partners of action clusters to excitatory neurons in cue clusters (*J*_CA,I_*J*_I*−*_*J*_EI_) by 40%. The full synaptic matrix for the whole model network is shown in **Supplementary Figure 4**.

### HMM analysis of the model simulations

For HMM analysis of model data, 10 different networks were used to mimic variability across sessions and/or subjects. The networks were obtained each time by sampling a new synaptic matrix (all having the same basic structure described earlier). Although each network had random connectivity, the in-degree connectivity from each presynaptic population was fixed across the 10 networks. This enforced some degree of uniformity in firing rates across clusters and minimized the presence of bias in the performance of each network.

A simulation of each network was run for 100 trials (25 trials per taste stimulus). For each trial, each neuron started with a random initial membrane potential (normally distributed with mean 0 and standard deviation 4) and received a constant external input current. At the start of the trial, the network was presented with one of the stimuli (which modified the external input current for select neurons) and evolved over 3 seconds. We then sampled one neuron per excitatory cluster, excluding those that fired at < 2 Hz on average whenever possible. An HMM was fitted to the spike trains of the sampled neurons following the same procedure used for the HMM analysis of experimental data. Trials were scored by inferring the model’s selected direction from the firing rates of its action clusters. Cluster firing rates were calculated by binning spikes in 50 ms windows, and a cluster was considered “on” when its firing rate was at least 40% of its maximum observed over the 100-trial simulation (setting a constant threshold at 34 Hz yielded exactly the same results). A trial was deemed correct if the appropriate action cluster came on after 0.5 s after stimulus onset while the opposite action cluster did not. A trial was deemed incorrect if the opposite action cluster came on during this window and the correct one did not. For an ambiguous trial, in which neither action cluster came on or both action clusters came on, we assigned a decision randomly such that the expected accuracy was 0.5, but also excluded the trial from further analysis. We then used the scored trials to classify the coding properties of the hidden states from each HMM, following the same procedure used for the experimental data. Onset times of all coding states were pooled over the 10 models, and distributions of these onset times were compared for Quality-coding states, Cue-coding states, and Action-coding states.

### Simulated optogenetic silencing experiments

The same 10 networks (characterized by the same 10 synaptic weight matrices used above) were subjected to various perturbations in a series of simulations meant to mimic experimental optogenetic silencing of cortical activity (Vincis et al., 2020). Silencing was implemented as a simple, square stimulus pulse to all inhibitory neurons in the network. The height of the square pulse quantified the percent increase in baseline external current to inhibitory neurons. As a function of the square pulse’s center *c*, its duration ∆_sil_, and its strength *κ*_sil_, the formula for the square pulse in time *t* read:

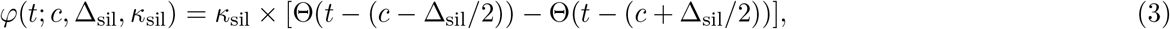

where Θ is the Heaviside step function. For silencing during the sampling period, *κ*_sil_ = 100%, ∆_sil_ = 500 ms, and *c* = 250 ms. For silencing during the delay period, *κ*_sil_ = 100%, ∆_sil_ = 2, 500 ms, and *c* = 1, 750 ms.

To test network performance across silencing conditions, we ran two independent ANOVAs, one for the stimulus with gain 60% and decay time constant 160 ms, and one for the stimulus with 60% and decay time constant 705 ms. These were 1-way within-subjects (repeated measures) ANOVAs with a single categorical factor, Silencing (levels: Full, Sampling, Delay). Both were significant and Bonferroni-corrected post-hoc tests were conducted (**Supplementary Figure 5a,c**). Network performances across different silencing and stimulus conditions for our best stimulus (gain 200% and decay time constant 160 ms) were compared using a 2-way within-subjects ANOVA with 3-level categorical factors, Silencing (levels: None, Sampling, Delay) and Stimulus (levels: Full, Head, Tail). The interaction was significant, and post-hoc analysis was conducted using a Bonferroni correction (**Figure 6b,d**).

We further tested perturbation strengths of *κ*_sil_ = 25% and *κ*_sil_ = 100% for a constant duration of ∆_sil_ = 250 ms and variable *c*. Here we computed mean network accuracy as a function of the silencing center *c* by “sliding” the square pulse center along the trial window from 0 to 3 s in increments of ∆*c* = 50 ms. For each value of *c*, we simulated 100 trials for each of the 10 networks under the given parameter set {*κ*_sil_, ∆_sil_}, and then averaged the accuracy over the collated 1, 000 trials.

For HMM analysis of a network with and without silencing perturbations, we chose the network whose average baseline accuracy was closest to the mean (samples shown in **Figure 6b, bottom left**). We reused the initial conditions from its 100 control trials 6 times, each time with a different silencing perturbation, either *κ*_sil_ = 25% (“Weak”) or *κ*_sil_ = 100% (“Strong”) and either *c* = 125 ms (“Beginning”), *c* = *t*^Cue^ (“Cue onset”), or *c* = 1, 500 ms (“Middle”), where 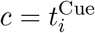 is the cue cluster onset time for the *i*t control trial, determined by thresholding the firing rates of the cue clusters as done for action clusters to infer network choice. In all cases ∆_sil_ = 250 ms. A single HMM was then fitted to this collection of 700 trials and used to decode, trial-by-trial, the sequence of hidden states shown in **Figure 7a**. Classification of the coding properties of these states (for **Figure 7c**) was done as before by comparing state occurrence frequencies across different stimulus and outcome conditions. In each trial, an Action-coding state is called “Correct” if its direction preference in correct trials matches the current trial’s correct direction (e.g., an Action-coding state with left direction preference in correct trials is considered “Correct” when it appears in Sucrose and Quinine trials, and is considered “Incorrect” when it appears in Maltose and Octaacetate trials).

To quantify hidden state sequence similarity over time (**Figure 7b**), we compared the average state sequences under conditions of silencing and control (i.e., in the absence of silencing) in the following way. We discretized time into 2 ms bins (full resolution of the HMM) and, *for each time bin*, obtained a vector ***x*** of 25 dimensions, one for each trial where the same taste stimulus was given (for example, sucrose). Each dimension contained either the current hidden HMM state in that trial, or a zero if no clear HMM state had been decoded in that trial. For example, if ***x*** = {3, 2, 0, …, 1}, in the first trial the HMM state was 3, in the second trial it was 2, in the third trial no state had been decoded in that bin, and so on, until the last trial, where state 1 was the decoded HMM state. A vector ***x***^*k*^ was built for each silencing condition *k* shown at the top of **Figure 7a**, and an analogous vector ***x***_0_ (in the same bin) was built for sequences under control (i.e., under no silencing). From each vector ***x***^*k*^ we built a measure of similarity with ***x***_0_ by counting how many times two states were the same across all 25 trials. This required summing up 25 × 25 comparisons 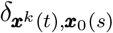, where *t, s* = 1, …, 25, index the trial (i.e., the elements of vectors ***x***^*k*^, ***x***_0_), and *δ*_*a,b*_ = 1 if *a* = *b* and *δ*_*a,b*_ = 0 otherwise. We performed this for all 4 taste stimuli and took the average. Since the total number of trials across all stimuli was 2, 500 = 25 × 25 × 4, the final measure of similarity in a single bin was

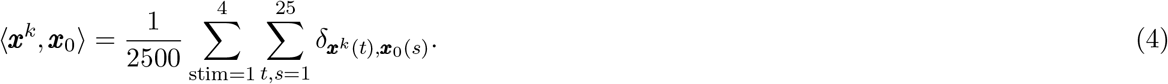

We finally normalized this measure by the similarity of *x*_0_ with itself:

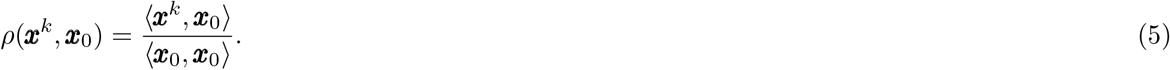

We repeated this procedure for all silencing conditions in **Figure 7a (top)**, i.e., for *k* standing for “Weak; Beginning”, “Strong; Beginning”, etc., and plotted the time course of each *ρ*(***x***^*k*^, ***x***_0_) in **Figure 7b** after convolving each curve with a Gaussian filter with binwidth of 150 time bins.

The hidden states shown in **Figure 7b** were classified for their coding properties by comparing their occurrence frequencies across trials based on stimulus and outcome (regardless of the silencing condition) using the same rules we followed previously. States classified as “Action-coding” were considered “Correct” or “Incorrect” based on their “preferred direction” in correct trials (see *Classification of decoded states*). If this direction matched the trial’s correct direction, the state was “Correct”; otherwise, it was “Incorrect.” Fractions of trials containing Correct Action-coding states were compared over all 7 conditions with a Chi-squared test. This was significant, so Marascuilo post-hoc tests were conducted to compare individual silencing conditions to the control condition (**Figure 7c**).

## Acknowledgements

We thank Dr. Luca Mazzucato, Xiaoyu Yang, and Tianshu Li for sharing their computer code, and Dr. Arianna Maffei for helpful discussions and feedback on the manuscript. This work was partially supported by a U01 grant from the NIH/NINDS Brain Initiative (1UF1NS115779) to G.L.C. and A.F and by grants R01DC015234 and R01DC018227 to A.F. The content of this article is solely the responsibility of the authors and does not necessarily represent the official views of the National Institutes of Health.

## Supplementary Information

**Supplementary Table 1.**
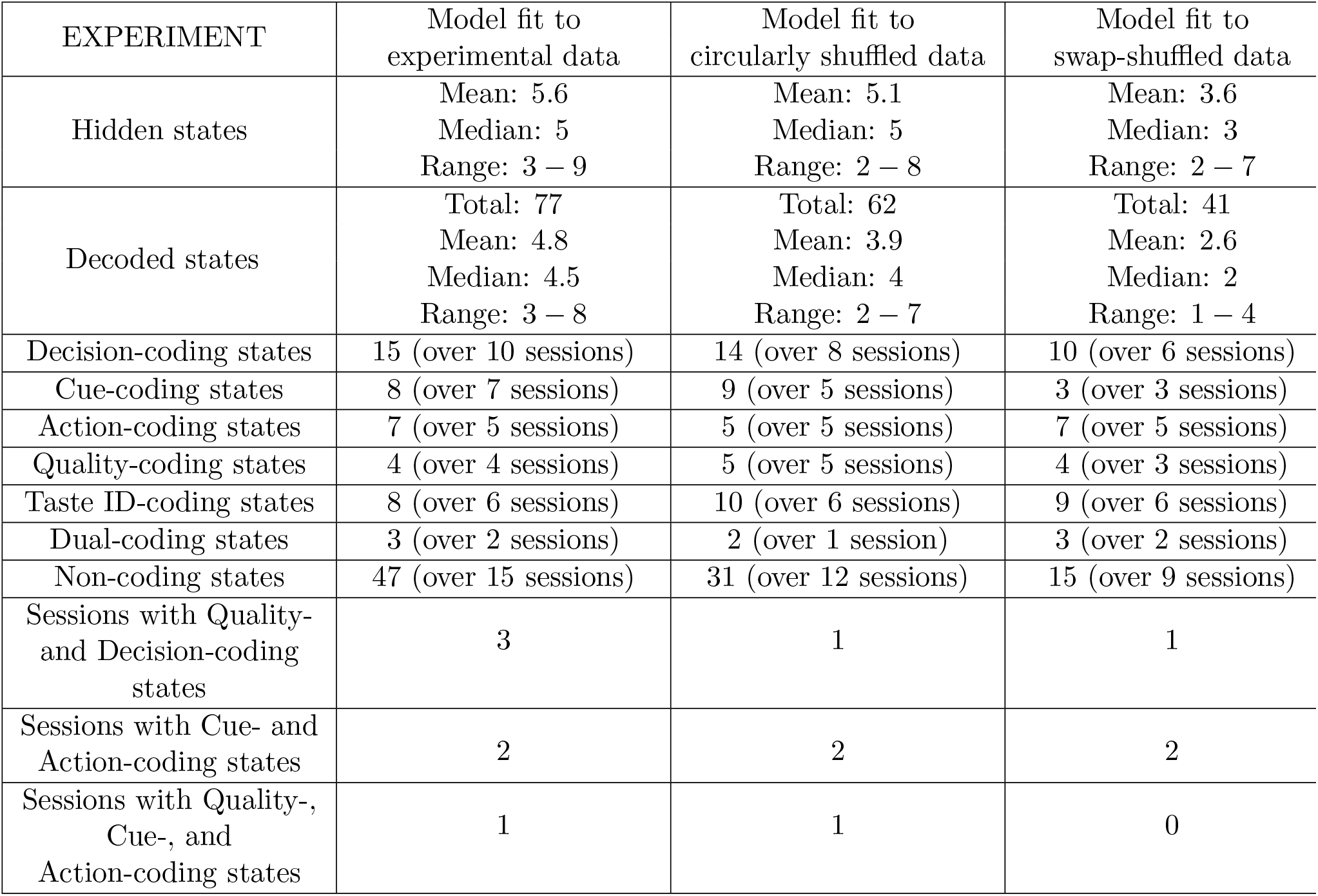
Summary of the numbers of states found by HMM models fit to unshuffled, circularly shuffled, and swap-shuffled experimental data. Also indicated is the distribution of these states over sessions (16 sessions total). The same coding state classification procedure (see. **Supplementary Figure 1**) was applied to the trial-by-trial decoding results from each model. “Hidden states” is the theoretical best number determined from model fitting; “Decoded states” is how many were actually found after decoding trial-by-trial.

**Supplementary Table 2.** Summary of numbers of states found by HMM models fit to unshuffled, circularly shuffled, and swap-shuffled simulation data. Also indicated is the distribution of these states over sessions (10 sessions total). The same coding state classification procedure (see **Supplementary Figure 1**) was applied to the trial-by-trial decoding results from each model. “Hidden states” is the theoretical best number determined from model fitting; “Decoded states” is how many were actually found after decoding trial-by-trial.

| SIMULATION | Model fit to<br>simulation data | Model fit to<br>circularly shuffled data | Model fit to<br>swap-shuffled data |
| --- | --- | --- | --- |
| Hidden states | Mean: 21.2<br>Median: 21<br>Range: 17 – 25 | Mean: 18<br>Median: 17<br>Range: 14 – 24 | Mean: 10.6<br>Median: 11<br>Range: 8 – 14 |
| Decoded states | Total: 174<br>Mean: 17.4<br>Median: 17<br>Range: 15 – 21 | Total: 158<br>Mean: 15.8<br>Median: 15.5<br>Range: 13 – 20 | Total: 81<br>Mean: 8.1<br>Median: 8<br>Range: 6 – 10 |
| Decision-coding states | 83 (over 10 sessions) | 74 (over 10 sessions) | 4 (over 3 sessions) |
| Cue-coding states | 50 (over 10 sessions) | 52 (over 10 sessions) | 1 |
| Action-coding states | 21 (over 10 sessions) | 15 (over 8 sessions) | 1 |
| Quality-coding states | 20 (over 10 sessions) | 2 (over 2 sessions) | 0 |
| Taste ID-coding states | 39 (over 10 sessions) | 49 (over 10 sessions) | 46 (over 10 sessions) |
| Dual-coding states | 0 | 0 | 1 |
| Non-coding states | 32 (over 10 sessions) | 33 (over 9 sessions) | 30 (over 10 sessions) |
| Sessions with Quality-<br>and Decision-coding<br>states | 10 | 2 | 0 |
| Sessions with Cue- and<br>Action-coding states | 10 | 8 | 0 |
| Sessions with Quality-,<br>Cue-, and<br>Action-coding states | 10 | 1 | 0 |

**Supplementary Table 3.**
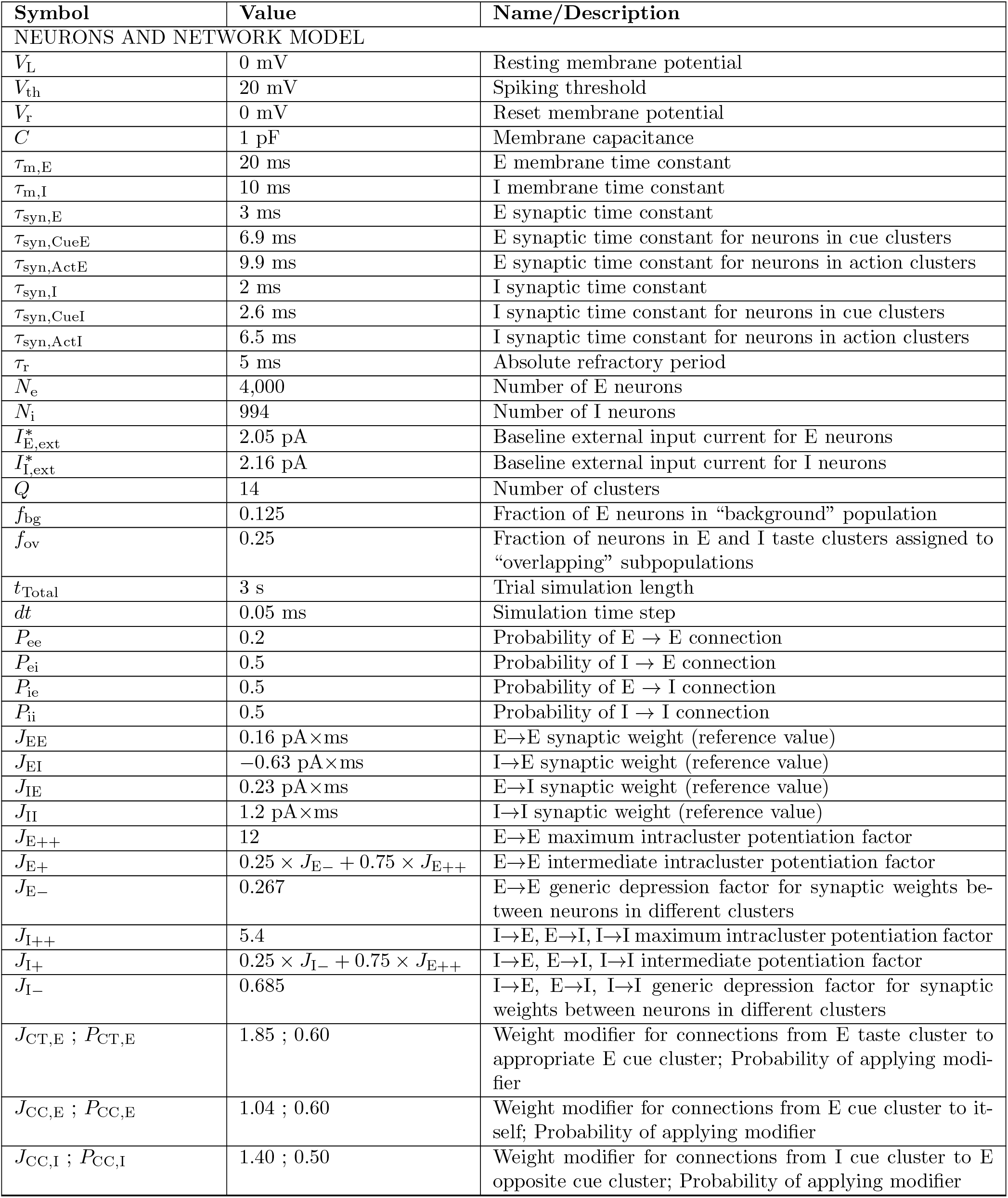
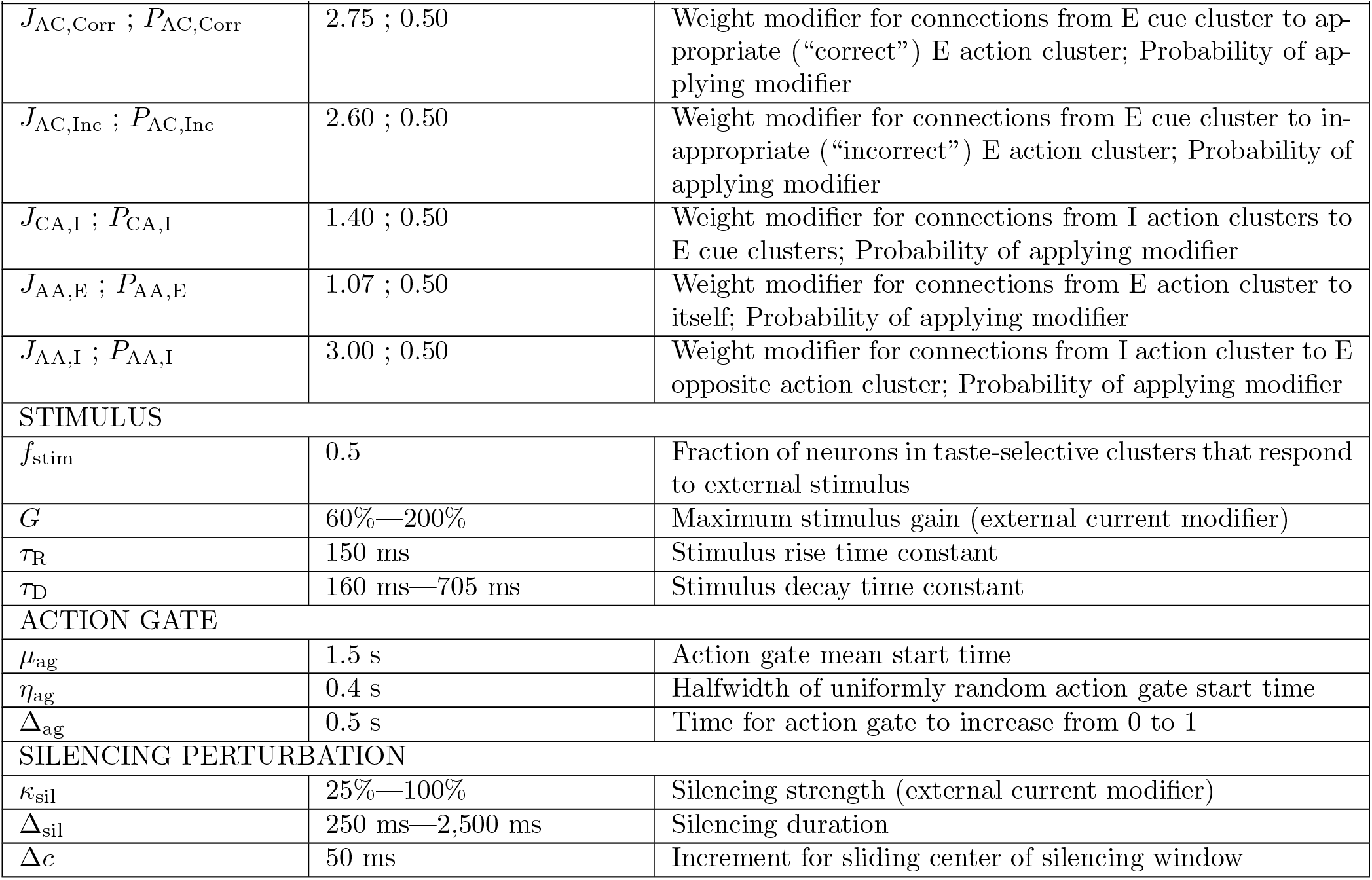
Model parameters. Symbols, values, and brief descriptions for all parameters used in the simulations of the spiking network model presented in the main text.

**Supplementary Figure 1.**
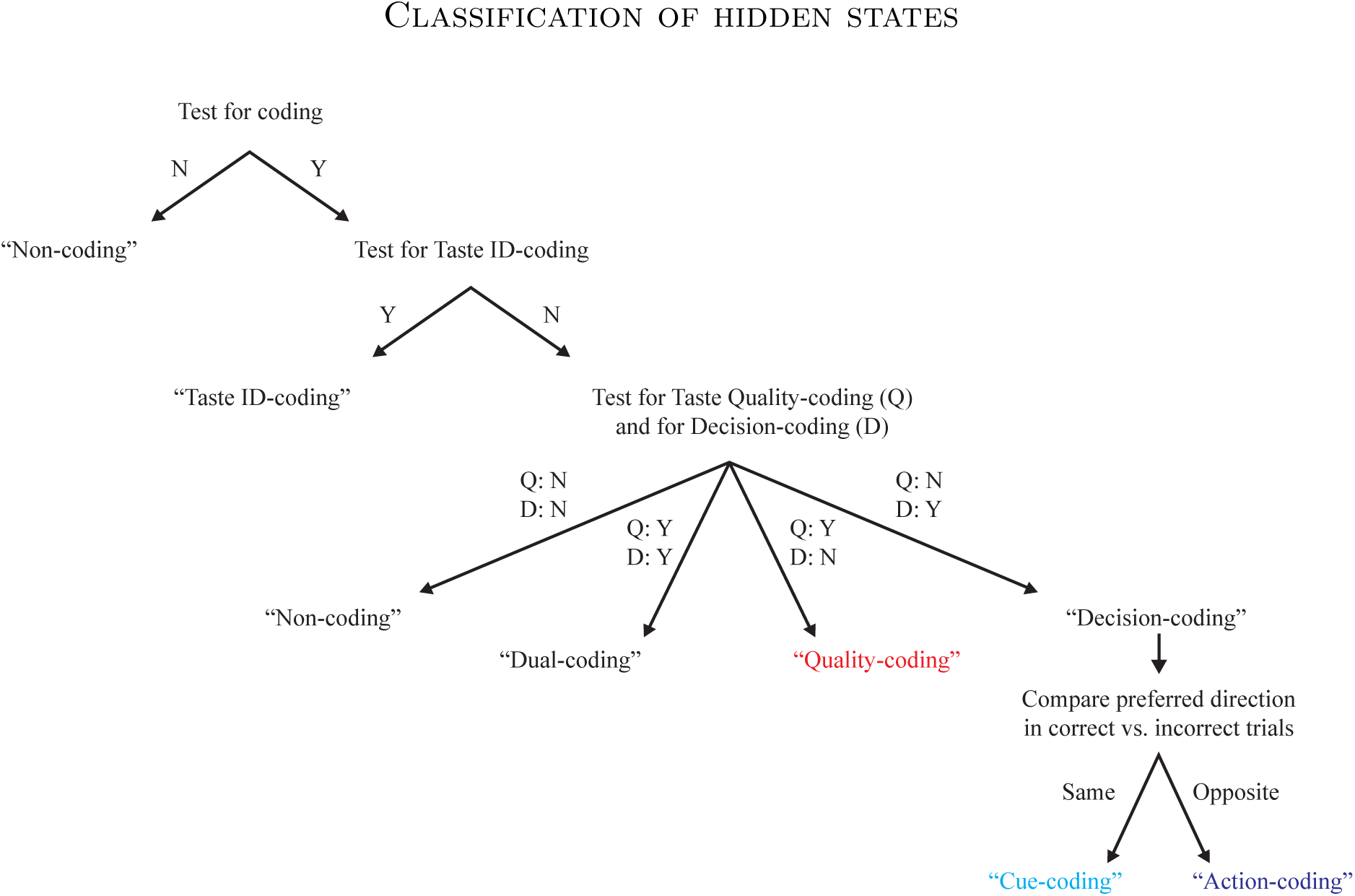
Pipeline for classification of HMM states. Given a state *S*, the probability of observing *S* in a correctly performed trial was compared across the four trial types using a Chi-squared test. If the result was not significant (N), *S* was labeled “Non-coding.” Otherwise (Y), all six pairwise Marascuilo post-hoc tests were performed. If there was exactly one trial type for which all post-hoc tests involving it were significant (Y), *S* was labeled “Taste ID-coding.” Otherwise (N), the four trial types were grouped in two orthogonal ways: sweet (sucrose and maltose) and bitter (quinine and sucrose octaacetate), and cue left (sucrose and quinine) and cue right (maltose and sucrose octaacetate). Two additional Chi-squared tests were run: one to compare the probability of observing *S* in correctly performed trials between the sweet and bitter categories, and another to compare the probability of observing *S* in correctly performed trials between the cue left and cue right categories. These two tests are run independently, leading to four possible significance outcomes. If neither test was significant, *S* was labeled “Non-coding.” If both tests were significant, *S* was labeled “Dual-coding.” If the sweet vs. bitter test was significant and the cue left vs. cue right test was not, *S* was labeled “Quality-coding.” If the sweet vs. bitter test was not significant and the cue left vs. cue right test was, *S* was labeled “Decision-coding,” but could be further classified by examining its frequency of occurrence in correctly vs. incorrectly performed trials. If the cued direction for which *S* was more likely to be observed in correctly performed trials was the same for incorrectly performed trials (referred to as the “preferred direction” for *S* in correct and incorrect trials), *S* was labeled “Cue-coding.” If the preferred directions in correct and incorrect trials were opposite, *S* was labeled “Action-coding.”

**Supplementary Figure 2.**
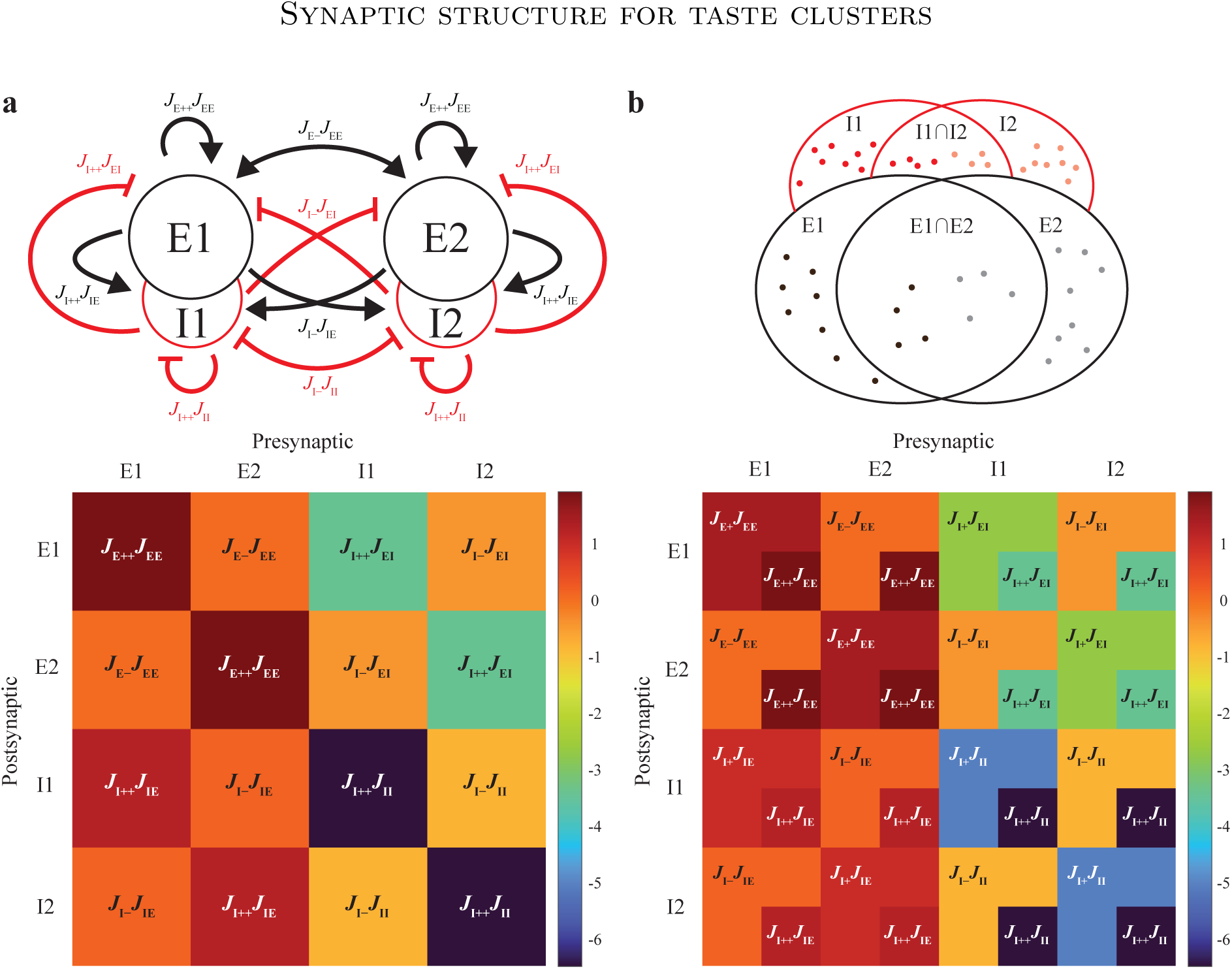
Synaptic structure for taste clusters. **a**: Generic synaptic structure between clusters without task roles. **Top**: Cartoon representing a pair of E-I partner clusters with their synaptic connection weights (red, flathead arrows: inhibitory connections; black, pointed arrows: excitatory connections). **Bottom**: synaptic matrix for the generic pair of E-I partner clusters (colormap corresponding to units of pA×ms). Given any two excitatory clusters, E1 and E2, and their inhibitory partners, I1 and I2, synaptic connections were strong between neurons within the same cluster or belonging to partner clusters (*J*_E++_*J*_EE_ or *J*_I++_*J*_*αβ*_, with *α, β* ∈ {E, I}), and weak between neurons in different clusters (*J*_E*−*_*J*_EE_ or *J*_*I−*_*J*_*αβ*_, with *α, β* ∈ {E, I}). **b**: **Top**: Venn diagram depicting the “overlapping” structure of two E-I taste cluster pairs with the same quality (sweet or bitter). **Bottom**: synaptic matrix of “overlapping” taste clusters depicted at top (colormap corresponding to units of pA×ms). Taste neurons in different clusters with the same taste quality had strongly connected overlapping subclusters (E1∩E2 and I1 I2 in the Venn diagram), with synaptic strength *J*_E++_*J*_EE_ or *J*_I++_*J*_*αβ*_, with *α, β* ∈ {E, I}. These overlapping subclusters are shown in the bottom right corner of each taste cluster in the synaptic matrix. Taste neurons in the same cluster but outside the overlapping subclusters were connected with synaptic strength *J*_E+_*J*_EE_ or *J*_I+_*J*_*αβ*_, with *α, β* ∈ {E, I}. All other neurons were connected as in the generic clusters of panel **a**, with *J*_E*−*_ < 1 *< J*_E+_ *< J*_E++_ and *J*_I*−*_ < 1 *< J*_I+_ *< J*_I++_.

**Supplementary Figure 3.**
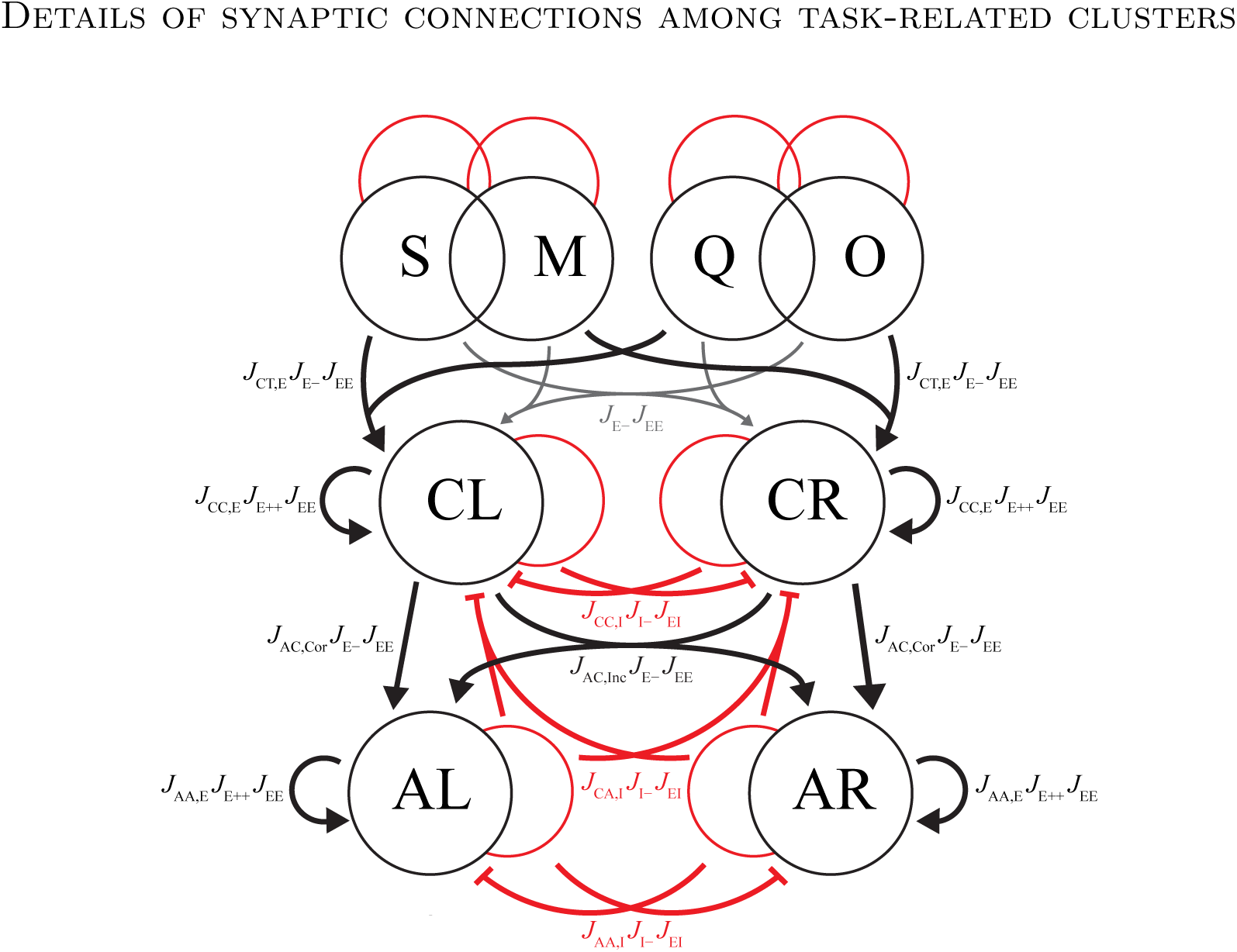
Details of synaptic connections among taste, cue, and action E-I cluster pairs (compare with main **Figure 3b**). The main synaptic connections are depicted by curved lines (pointed arrowheads are excitatory connections, flat arrowheads are inhibitory connections). The strength of each connection is indicated next to the corresponding curved line (see **Supplementary Table 3** for parameter values). Synaptic weights from excitatory taste clusters to appropriate excitatory cue clusters (i.e., S → CL, M → CR, Q → CL, and O → CR) were larger by a factor *J*_CT,E_ compared to generic weights between different clusters (e.g., S → CR, M → CL, Q → CR, and O → CL). Excitatory cue clusters had stronger intracluster connections (by a factor *J*_CC,E_) compared to generic clusters; the synaptic weights from an inhibitory cue cluster to the opposite excitatory cue cluster were larger by a factor *J*_CC,I_. Connections from excitatory cue clusters to correct excitatory action clusters (i.e., CL → AL and CR → AR) were larger by a factor *J*_AC,Cor_, while the connections to incorrect excitatory action clusters (i.e., CL → AR and CR → AL) were larger by a factor *J*_AC,Inc_, with *J*_AC,Cor_ *> J*_AC,Inc_. Excitatory action clusters had stronger intracluster connections (scaled up by *J*_AA,E_) and the weights from their inhibitory partners to the opposite excitatory action cluster were scaled up by *J*_AA,I_. Connections from inhibitory action clusters to excitatory cue clusters were magnified by *J*_CA,I_. All synaptic weight modifiers were applied with a given probability (reported in **Supplementary Table 3**). Connections among clusters not explicitly depicted here were as in **Supplementary Figure 2** (also see **Supplementary Figure 4** for the full synaptic weight matrix). Key: S: sucrose, M: maltose, Q: quinine, O: octaacetate, CL: cue left, CR: cue right, AL: action left, AR: action right.

**Supplementary Figure 4.**
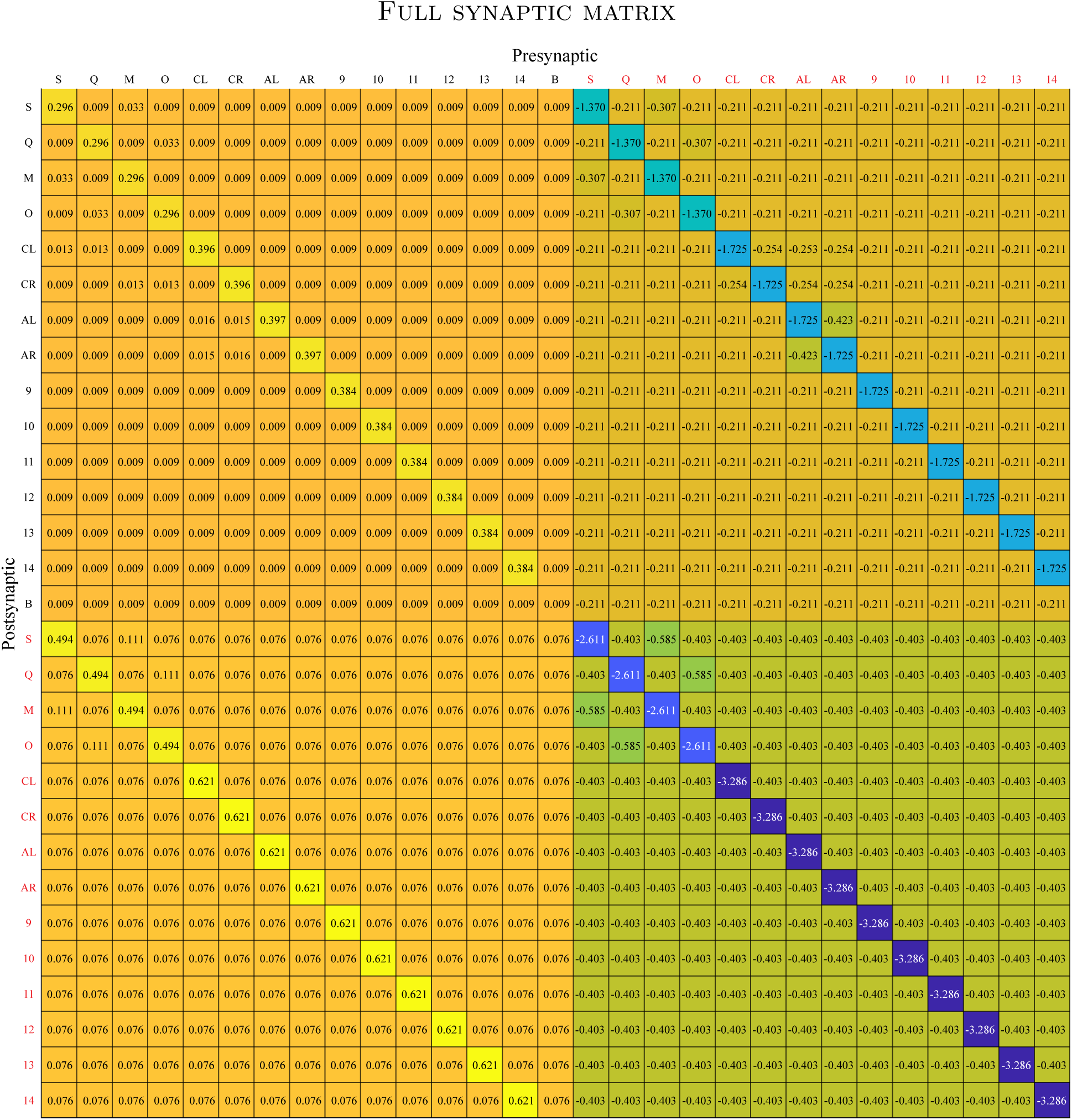
Full synaptic weight matrix of the network model. The value in a block occupying row *i* and column *j* is the *average* synaptic weight (in units of pA×ms) from a presynaptic neuron belonging to cluster *j* to a postsynaptic neuron belonging to cluster *i*. The average synaptic values shown in the table were obtained (u to rounding) as: 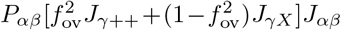 for connections involving taste clusters with the same taste quality (sweet or bitter), and 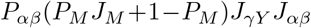 for all other connections. Here *α, β, γ* ∈ {E, I}, *γ* = E if and only if *α* = *β* = E, *X* ∈ {−, +}, *Y* ∈ {−, ++}, *J*_*M*_ ∈ {*J*_CT,E_, *J*_CC,E_, *J*_CC,I_, *J*_AC,Cor_, *J*_AC,Inc_, *J*_AA,E_, *J*_AA,I_, *J*_CA,I_} is a weight modifier, and *P*_*M*_ ∈ {0, *P*_CT,E_, *P*_CC,E_, *P*_CC,I_, *P*_AC,Cor_, *P*_AC,Inc_, *P*_AA,E_, *P*_AA,I_, *P*_CA,I_} is the corresponding probability of applying the modifier (*P*_*M*_ = 0 when modifier was not applied). Key: S: sucrose cluster, M: maltose cluster, Q: quinine cluster, O: octaacetate cluster, CL: cue left cluster, CR: cue right cluster, AL: action left cluster, AR: action right cluster, 9-14: clusters without task roles, B: background (excitatory) population

**Supplementary Figure 5.**
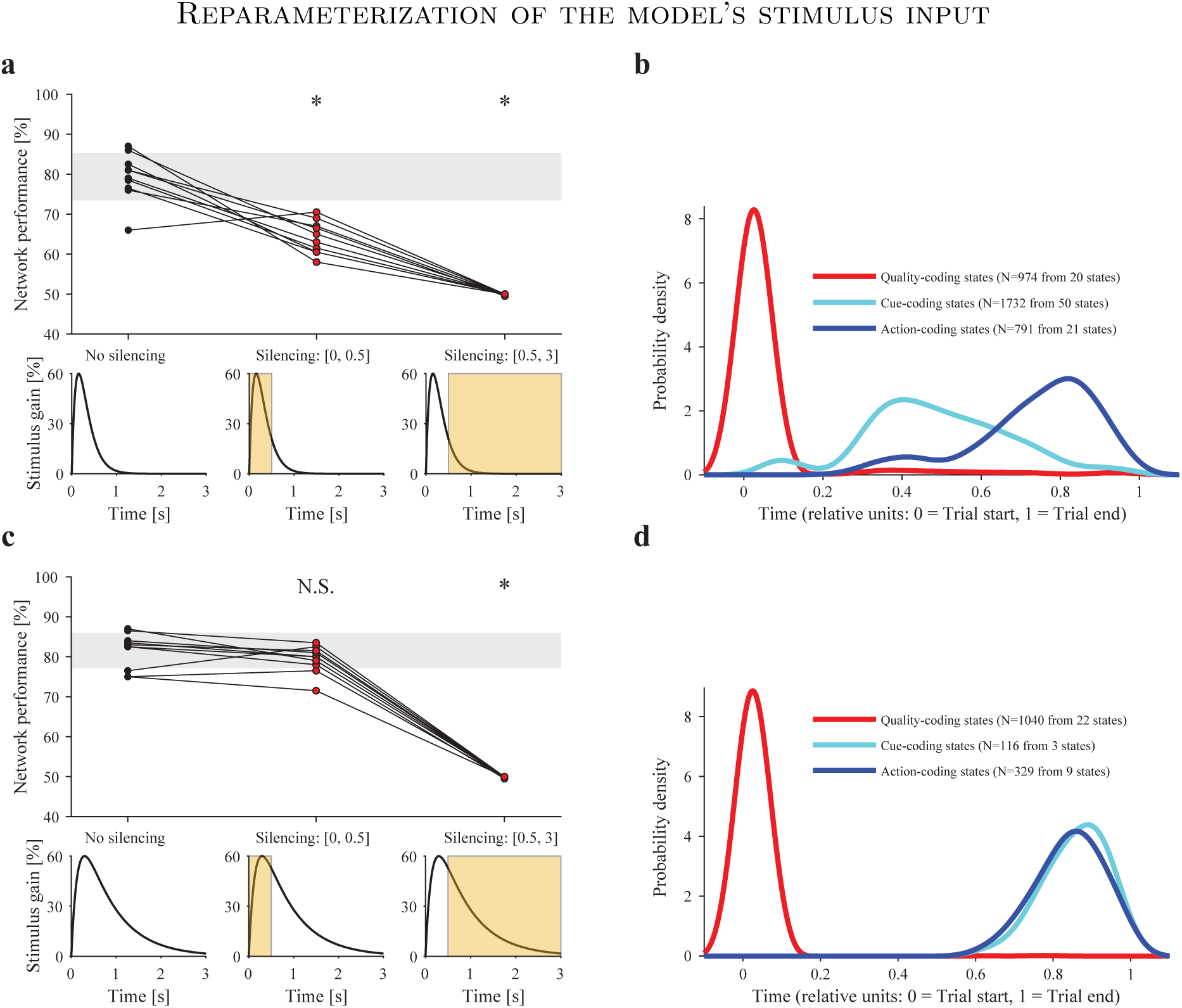
Network’s performance and onset times of coding states for different parameters of the stimulus input (supplements main **Figure 6**). **a, c**: Effect of simulated silencing during sampling and delay periods on task performance for models with stimulus input with gain 60% and decay time constant 160 ms (**a**) and 705 ms (**c**), respectively. **b, d**: Distribution of onset times of coding states after fitting HMMs to models with stimulus input as in corresponding left panel. * indicates significant difference (*p* < 0.05) for Bonferroni-corrected post-hoc test vs. None condition after significant within-subjects ANOVA across the 3 conditions. N.S. indicates no significant difference.

**Supplementary Figure 6.**
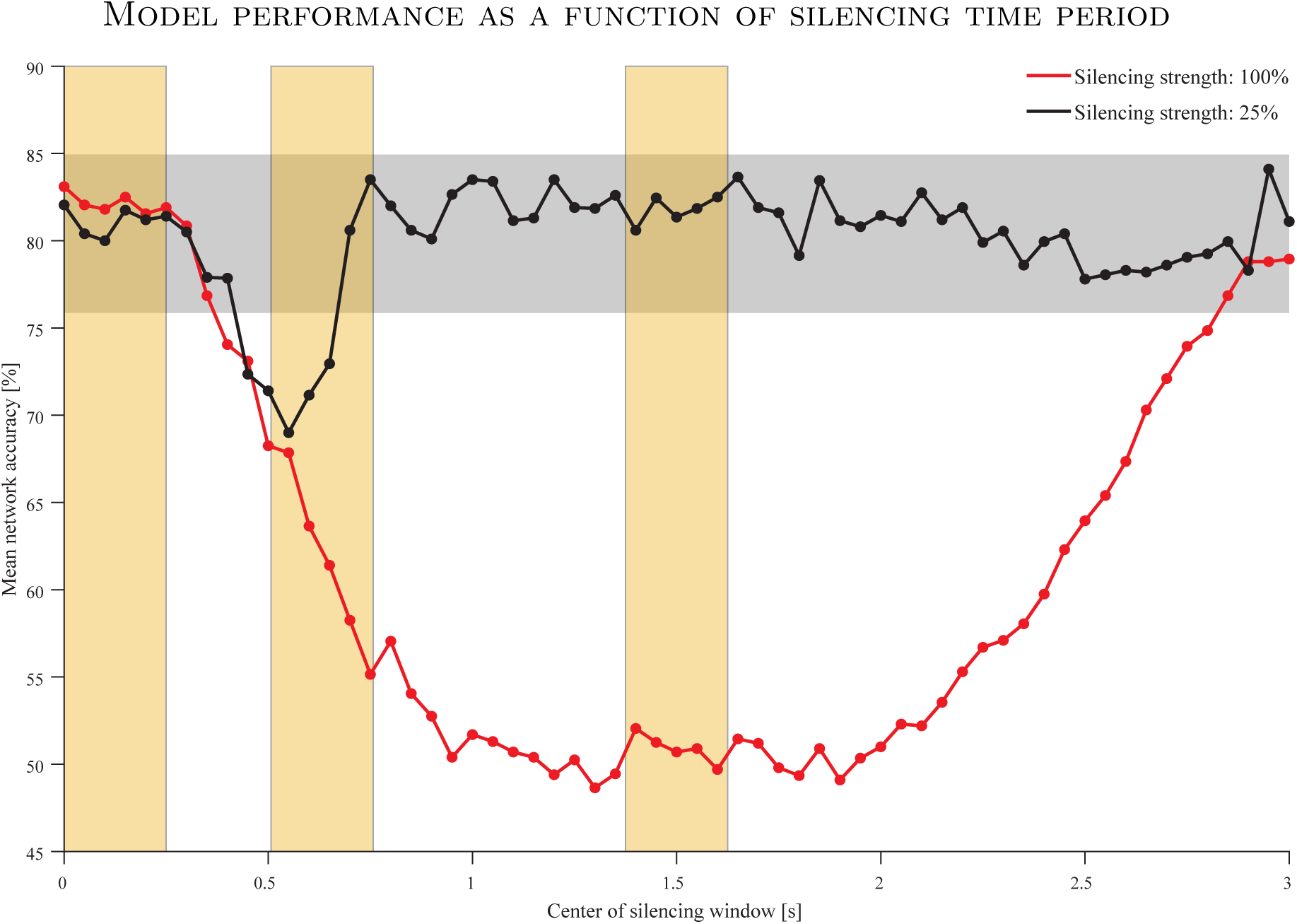
Effect of simulated optogenetic silencing on network task performance as a function of time of silencing. Silencing was implemented as a square pulse stimulus with width 250 ms and height (strength, 100% or 25%) determining the increase in baseline external current for all inhibitory neurons in the network. Each point is an average accuracy over 1, 000 trials (100 trials from each of 10 networks) for silencing centered at that point. For all points, the input stimulus had a gain of 200% and decay time constant of 160 ms. Grey region represents the mean ±1 standard deviation of accuracy for networks with no silencing (same as in main **Figure 6b**). Regions-of-interest for silencing employed in main **Figure 7a** (Beginning, Cue onset, and Middle) are shaded in yellow.

